# Whole-brain dynamical modeling for classification of Parkinson’s disease

**DOI:** 10.1101/2022.06.08.495360

**Authors:** Kyesam Jung, Esther Florin, Kaustubh R. Patil, Julian Caspers, Christian Rubbert, Simon B. Eickhoff, Oleksandr V. Popovych

## Abstract

Simulated whole-brain connectomes demonstrate an enhanced inter-individual variability depending on data processing and modeling approach. By considering the human brain connectome as an individualized attribute, we investigate how empirical and simulated whole-brain connectome-derived features can be utilized to classify patients with Parkinson’s disease against healthy controls in light of varying data processing and model validation. To this end, we applied simulated blood oxygenation level-dependent signals derived by a whole-brain dynamical model simulating electrical signals of neuronal populations to reveal differences between patients and controls. In addition to the widely used model validation via fitting the dynamical model to empirical neuroimaging data, we invented a model validation against behavioral data, such as subject classes, which we refer to as behavioral model fitting and show that it can be beneficial for Parkinsonian patient classification. Furthermore, the results of machine-learning reported in this study also demonstrated that performance of the patient classification can be improved when the empirical data are complemented by the simulation results. We also showed that temporal filtering of blood oxygenation level-dependent signals influences the prediction results, where the filtering in the low-frequency band is advisable for Parkinsonian patient classification. In addition, composing the feature space of empirical and simulated data from multiple brain parcellation schemes provided complementary features that improve prediction performance. Based on our findings, we suggest including the simulation results with empirical data is effective for inter-individual research and its clinical application.

## Introduction

For decades, large-scale whole-brain connectivity acquired from non-invasive *in-vivo* MRI has actively been used to study the human brain as an integrative complex system.^1^ Accordingly, anatomical (or structural) and functional connectivities between brain regions have been used. Previous studies have shown that the structural architecture shapes temporal synchronization between the blood oxygenation level-dependent (BOLD) signals in selected networks, for instance the default mode network.^2,3^ However, the structure-function correspondence is not high for whole-brain connectivity.^4-6^ The correspondences between the brain connectomes from the same and different subjects, samples or data modalities^7,8^ have been considered to investigate the inter-individual differences^9^ or diagnostic classification between healthy controls (HC) and patients.^4,10-12^

Connectivity relationships are also commonly used when the brain dynamics are modeled by mathematical whole-brain dynamical models. In particular, finding the strongest correspondence (the highest similarity) between empirical functional connectivity (eFC) and simulated functional connectivity (sFC) has been used for model validation.^13-15^ Such a correspondence of the simulated data to the empirical data may undergo qualitative changes when parameters of a given model vary, and the validation procedure consists in finding the most pronounced agreement between the data and the model fitted by searching for optimal parameter points.

Previous studies utilizing the discussed whole-brain modeling showed that the employed modeling approach is applicable to clinical research. Variability of the model parameters between diseased and healthy states has been investigated for brain disorders including schizophrenia,^16-19^ Alzheimer’s disease,^20^ Parkinson’s disease (PD),^21,22^ and stroke patients.^23^ For instance, Saenger *et al*.^22^ showed that therapeutic deep brain stimulation in PD can be modeled by the normal form of a Hopf bifurcation model.^24^ Detailed simulations of neuronal dynamics may also provide a way to test prognostic outcomes *in silico* throughout virtual operations and optimize setups and parameters of therapeutic interventions.^25-28^

There are, however, no well-established standards for model validation against empirical data. Several fitting modalities have been suggested in the literature including the fitting of the grand-averaged empirical and simulated FC matrices, fitting the dynamical FCs, maximization of the metastability, and structure-functional model fitting.^6,13,24,29,30^ On that account, it is necessary to investigate which parameter points of a given dynamical mode and which model fitting modalities are the most suitable to answer a given research question by the modeling approach. For example, it was observed that distributions of the optimal model parameters differ when using only functional or structure-functional model fitting and may lead to subject stratifications showing different model fitting values and optimal parameter points.^30^ It is also well known that varying parameters of MRI data processing influence the empirical structural and functional connectomes and their analyses.^31-34^ This subsequently affects model validation.^6,30,35^ Therefore, the impact of data processing on the results of model validation should carefully be considered, especially in clinical applications.

In PD research, eFC of the resting-state networks was already used in machine-learning approaches of subject classification.^36,37^ When sFC is involved, it is essential to extract relevant features for PD classification from simulation results via searching a given model parameter space for the optimal model. To do this, we considered two aspects of parameters regarding dynamical models and data processing. First, we find the model parameters that reveal the most prominent differences in connectome-correspondence between PD and HC. Such an approach can be used for model validation. Here, we aim at a diagnostic classification of patients from healthy subjects, where the model fitting to empirical neuroimaging data may be a suboptimal choice, and a behavioral (phenotypical) fitting might be better. For instance, disease status of the subjects can be used for behavioral fitting as we show in this study. Second, we consider different temporal filters of BOLD signals, which are known to influence FC properties.^38,39^ In particular, the altered frequency bands were found to retain PD-related neural changes.^40^ The frequencies of empirical BOLD signals, when included in the whole-brain mathematical models, may influence the optimal model parameters and the quality of the model fitting.^6,30^ In this context, investigation of the impact of temporal filtering conditions on the model validation in PD data is important. In the current study we advance the classification of clinical data by application of machine learning to empirical and simulated connectomes. The functional connectomes were calculated from empirical and simulated BOLD signals, respectively, filtered in broad-, low- and high-frequency bands for two different brain parcellations as given by the Schaefer^41^ and the Desikan-Killiany^42^ brain atlases. As compared to purely empirical studies, we make a next step based on the two aspects of parameters for model fitting modality and data processing and employ the simulated data to improve the prediction results in a machine-learning setting.

The current study employs whole-brain dynamical modeling of the resting-state functional MRI data based on the Jansen-Rit model type of interacting excitatory and inhibitory neuronal populations.^43,44^ The simulated FCs generated for the optimal model parameters based on model fitting modalities were used to calculate the connectome relationships (Pearson’s correlation) with empirical structural and functional connectivities. We also introduced a simple but effective method for model validation against behavioral data more suitable for differentiation between patients with PD patients and HCs than the conventionally used model fit to neuroimaging data. Consequently, the personalized features derived from the connectome relationships were used in this study for classification of PD and HC using machine-learning. We, in particular, show that complementing empirical data with simulated FC can improve the prediction performance for unseen subjects. Our results suggest that the personalized whole-brain models can serve as an additional source of information relevant for disease diagnosis and possibly for their treatment as well.

## Materials and methods

We performed three main steps to obtain the whole-brain connectivities eFC, eSC (empirical streamline counts), ePL (empirical average path-length), and sFC. Fig. 1 schematically illustrates the data processing and simulation workflow. We applied four temporal filtering conditions to empirical and simulated resting-state BOLD signals. Subsequently, we considered three types of connectivity relationships corresponding to the correlation between eFC and eSC, the correlation between eSC and sFC, and the correlation between eFC and sFC. Due to sFC induced by varying the two free model parameters of global coupling and global delay, the correlations involving sFC change as illustrated by the eFC-sFC correlation landscape in the parameter space in Fig. 1 (the rightmost color plot). We used these three connectivity relationships as features for the PD classification via a machine-learning approach. To this end, we trained PD classifiers and evaluated their performance based on prediction probabilities obtained on unseen subjects.

**Figure 1.**
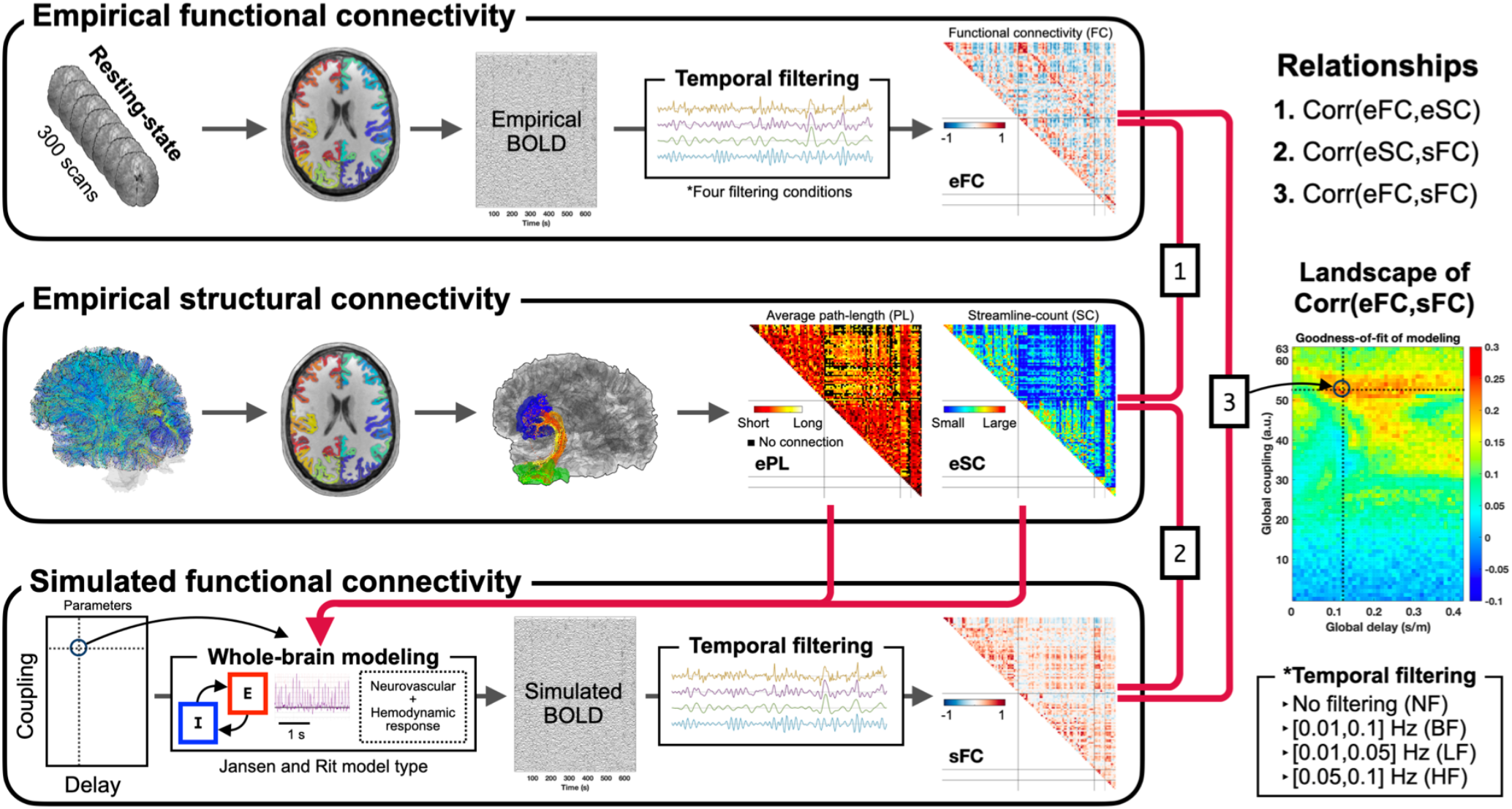
Data processing and simulation overview. First (upper box), brain parcellations in the native space of T1w were prepared and applied to the processed functional MRI data, BOLD signals were extracted from the corresponding brain regions and filtered according to four temporal filtering conditions (right bottom box), and four respective empirical functional connectivities (eFC) were calculated. Second (middle box), the parcellations were also used for calculation of the structural connectivity by extracting streamlines from the whole-brain tractography reconstructed using diffusion-weighted images, where the number and length of streamlines connecting any two brain regions were collected into matrices of empirical streamline count (eSC) and averaged path-length (ePL). Third (lower box), the structural connectome (eSC and ePL) was used to build a brain network for the whole-brain modeling that simulates BOLD signals, which were filtered according to the considered filtering conditions (right bottom box) and used to calculate simulated FC (sFC). Subsequently, we calculated connectivity relationships (Pearson’s correlation) using these three connectivity matrices: (1) corr(eFC, eSC), (2) corr(sFC, eSC), and (3) corr(eFC, sFC). Model parameters of global coupling and global delay were varied to validate the model against empirical data. In particular, the correspondence (correlation) between empirical connectivities (eSC and eFC) and simulated functional connectivities (sFC) was calculated for each parameter point resulting in the similarity landscapes in the model parameter space, see the example of the relationship between eFC and sFC in rightmost color plot. The most pronounced correspondence (correlation) between the empirical and simulated connectomes was selected together with the respective optimal model parameters as a result of the neuroimaging model fitting for further analysis.

### Subjects and demography

The three considered whole-brain connectivities (eFC, eSC, and sFC) were calculated for 51 (30 males) healthy controls and 65 (45 males) patients with Parkinson’s disease, see Table 1 for the demography. Patients and controls were included from an MRI data pool acquired at the University Hospital Düsseldorf, which was also used in several recent studies^36,37,45,46^ where additional details about the data can be found. All patients were diagnosed with PD by an experienced movement disorder specialist. All HC subjects had no history of any neurological or psychiatric disease and no abnormalities detected in cranial MRI. The ages of 116 subjects (mean: 58.9 years and standard deviation: 10.3 years) are in a normal distribution (the null hypothesis was not rejected by a Chi-square goodness-of-fit test with *p* = 0.15). The age of patients was significantly higher than that of controls (Wilcoxon rank-sum two-tail test). The age of male patients was significantly higher than that of male controls, but the age of females was not from distributions with different medians. There was no age difference between females and males (Table 1). The study was approved by the local ethics committee and performed in accordance with the declaration of Helsinki. All subjects provided written informed consent prior to study inclusion.

**Table 1.** Demography of subjects included in the study.

| Groups | Mean (standard deviation) years |  | Statistical tests | p-values |
| --- | --- | --- | --- | --- |
|  | **All subjects** |  | **Chi-square goodness-of-fit test** |  |
| All | 58.93 (10.25) |  | 116 subjects | 0.149 |
|  | **Healthy controls** | **Patients** | **Wilcoxon rank-sum two-tail test** |  |
| All | 55.02 (9.69) | 62.00 (9.62) | 51 healthy controls versus 65 patients | 0.000 |
| Female | 56.52 (9.40) | 60.80 (8.96) | 21 healthy controls versus 20 patients | 0.201 |
| Male | 53.97 (9.74) | 62.53 (9.85) | 30 healthy controls versus 45 patients | 0.001 |
|  | **Females** | **Males** | **Wilcoxon rank-sum two-tail test** |  |
| All | 58.61 (9.43) | 59.11 (10.67) | 41 females versus 75 males | 0.751 |

### MRI acquisition

Structural and functional MRI data were acquired using a 3T scanner (Siemens Trio). A structural brain image was acquired using a three-dimensional T1-weighted image (T1w) sequence (TR = 2.3 s, TE = 2.96 ms, TI = 900 ms, flip-angle = 9°, field-of-view = 240 × 256 mm^2^ in sagittal, the number of slices = 160, voxel dimension = 240 × 256 × 160, voxel size = 1.0 × 1.0 × 1.1 mm^3^). Diffusion-weighted images (DWI) comprised a single non-weighted (B0) image and weighted (B = 1000 s/mm^2^) images with 64 directions (TR = 6.7 s, TE = 81 ms, phase encoding: anterior to posterior, field-of-view = 216 × 216 mm^2^ in axial, the number of slices = 55, voxel dimension = 90 × 90 × 55, voxel size = 2.4 × 2.4 × 2.4 mm^3^). Resting-state functional MRI was obtained using an echo-planar imaging sequence during 663 s (TR = 2.21 s, TE = 30 ms, field-of-view = 200 × 200 mm^2^ in axial, the number of slices = 36, voxel dimension = 64 × 64 × 36, voxel size = 3.125 × 3.125 × 3.565 mm^3^). To prevent distraction of streamline tracking, artifact volumes of DWI were removed from the data based on evaluation by two raters.

### Preprocessing of MRI

For the personalized data processing, we developed a containerized in-house pipeline to process structural and functional MRI in the native spaces. The pipeline consists of five modules: preprocessing of structural MRI (T1w and DWI), whole-brain tractography (WBT) calculation, atlas transformation, reconstruction of structural connectivity (eSC and ePL), and preprocessing of functional MRI. The pipeline comprises Freesurfer,^47^ FSL,^48^ ANTs,^49^ MRtrix3,^50^ and AFNI.^51^ It is publicly available (https://jugit.fz-juelich.de/inm7/public/vbc-mri-pipeline).

The preprocessing module of structural MRI performed the following steps: bias-field correction for T1w, alignment of anterior-posterior commissures of T1w, *recon-all* by Freesurfer, removing the Gibbs ringing artifacts of DWIs, bias-field correction for DWIs, corrections of head motion, b-vector rotations and eddy distortion of DWIs, and co-registration between averaged DWI and T1w. This module segmented subcortical areas based on voxel intensities of the T1w images. It also prepared labeling annotations using a brain atlas for which a classifier was available from the literature. The annotation can also be created based on a subject cohort by capturing region data either drawn by neuroanatomists or according to dedicated algorithms.^52^

The WBT calculation module included only MRtrix3 functions. They estimated response functions for spherical deconvolution using the constrained deconvolution algorithm.^53^ Fiber oriented distributions (FOD) were estimated from the DWIs using spherical deconvolution, and the WBT was created through the fiber tracking by the second-order integration over the FOD by a probabilistic algorithm.^54^ In the latter step, we used 10 million total streamlines for the WBT density. The tracking parameters of *tckgen* function were set as in the previous study^30^: step size = 0.625 mm, angle = 45 degrees, minimal length = 2.5 mm, maximal length = 250 mm, FOD amplitude for terminating tract = 0.06, maximum attempts per seed = 50, maximum number of sampling trials = 1000, and downsampling = 3 (FOD samples per steps – 1).

The atlas transformation module annotated labels using a classifier to parcel cortical regions in the native T1w space using Freesurfer. In the present study, we applied two atlas classifiers for brain parcellations, the Schaefer atlas with 100 parcels^41^ and the Desikan-Killiany atlas with 68 parcels.^42^ Both atlases provide cortical parcellations, where the former is based on functional MRI data, while the latter is labeled by gyral-based anatomical parcellation. After this, the subcortical areas segmented by the preprocessing module were included and combined with the labeled cortical parcels. Finally, the pipeline transformed the labeled image (cortical parcels and subcortical regions) from the T1w to DWI native spaces.

The reconstruction module calculated the matrices of the streamline counts (SC) and the matrices of the average path lengths (PL) of the streamlines extracted between any two parcellated brain regions from the calculated WBT with the transformed labeled image in the DWI space.

The preprocessing module of functional MRI performed slice time correction, head motion correction, re-slicing in a 2 mm iso-cubic voxel space, intensity normalization, de-trending with filtering of very slow fluctuations out (high pass), co-registration to the T1w image, and calculation of regressors for the white-matter, CSF, and brain global signals as well as for the head motion. The pipeline also transformed the labeled image of the brain parcellation generated in the native T1w space to the functional MRI native space. Finally, we performed a nuisance regression with the prepared regressors (white-matter, CSF, and the brain global signals as well as head motions).

### Post-processing of functional MRI

After preprocessing of MRI, we extracted mean BOLD signals based on the annotated atlas labels and applied three temporal band-pass filtering conditions in the frequency ranges [0.01,0.1] Hz (broad-frequency band; BF), [0.01,0.05] Hz (low-frequency band; LF), and [0.05,0.1] Hz (high-frequency band; HF). Therefore, four filtering conditions were considered: no filtering (NF), BF, LF, and HF. The filtering was done using a script in the Python programming language (version 3.8, Python Software Foundation, https://www.python.org/) using the SciPy (version 1.5) signal processing module^55^ and NumPy^56^ (version 1.19) for the temporal band-pass filtering. We used the Butterworth digital filter of the order 6, *scipy*.*signal*.*butter*.

### Whole-brain model

#### Convolution-based two-population model for electrical signals

The whole-brain resting-state dynamics considered in this study was simulated by a system of N coupled neuronal models representing the mean brain regional activity. Each region contains two populations for each neuronal type (excitatory and inhibitory) that interact with each other via post-synaptic potentials (PSP).^43^ The considered convolution-based model is of the Jansen-Rit type^44,57^ and simulates the PSP signals involving other brain regions that interact with time delay in coupling according to the calculated structural connectivity, i.e., SC and PL matrices. The following set of differential equations describes the mean dynamics of the excitatory and inhibitory PSPs of region *n* = 1, 2, …, *N*:

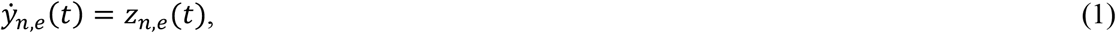

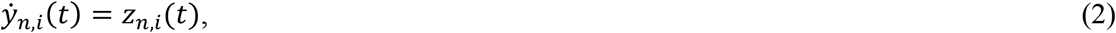

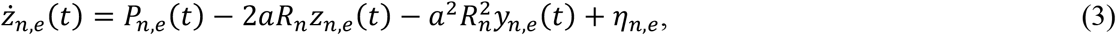

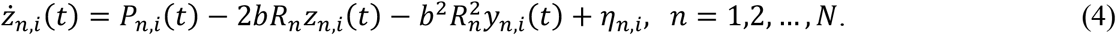

Here, *z*_*n,e*_, *z*_*n,i*_, *y*_*n,e*_, and *y*_*ni*_ are the excitatory post-synaptic current, the inhibitory post-synaptic current, the excitatory PSP (EPSP), and the inhibitory PSP (IPSP) of the brain region *n*, respectively, where the subscripts *e* and *i* stand for *excitatory* and *inhibitory*, accordingly. The model (1)-(4) is a system of driven harmonic oscillators in a critical damping regime, where the system quickly returns to its steady state after perturbation without under-shooting. Parameters *a* and *b* represent the reciprocal of the time constants of the PSP kernel for the two populations for EPSP and IPSP, respectively. *η*_*n,e*_ and *η*_*n,i*_ are independent noise sampled from a random uniform distribution between -1.5 and 1.5 *V/s*^*2*^. For frequency of oscillations, we also introduced a scaling factor *R*. By increasing *R*, the spectral power of the PSP signals shifts to higher frequencies. Perturbation *P*_*n,e*_ drives EPSP oscillations regarding input signals from other regions, i.e., it models the coupling between the network nodes/brain regions, and *P*_*n,i*_ perturbs IPSP oscillations by the input from the excitatory population in the same region *n*,

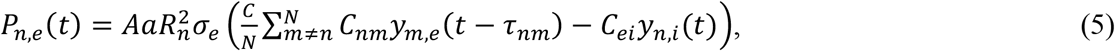

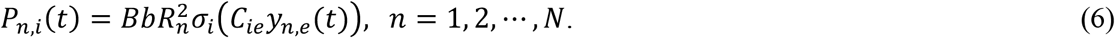

*A* and *B* are the maximum amplitudes of the PSP kernels for EPSP and IPSP, respectively. *N* is the total number of brain regions/network nodes for the whole-brain model. In Equation 5, *C* is a global coupling parameter which scales the couplings throughout the whole-brain network. *C*_*nm*_ is the strength of the individual coupling from region *m* to region *n*, which is realized via weighting the EPSP signal of the m-*th* network node *y*_*m,e*_ considered with time delay *τ*_*nm*_. Parameter *C*_*ei*_ weights an input coming from the inhibitory population of the same brain region, i.e., IPSP *y*_*n,i*_. The individual time delays and coupling strengths between regions *m* and *n* can be estimated from the empirical data as

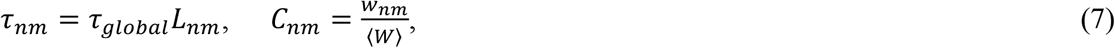

where the averaged path length *L*_*nm*_ (from the matrix PL) of the reconstructed streamlines between regions *n* and *m* is scaled by a global delay parameter *τ*_*global*_. *C*_*nm*_ in Equations 7 calculates an individual coupling strength by taking into account the SC matrix, where the number of streamlines *w*_*nm*_ between the two regions was normalized by an averaged number of streamlines ⟨*W*⟩ calculated over all connections except for the self-connections. As follows from Equation 5, the coupling between brain regions is realized between the excitatory populations, where the delayed EPSP signals from the other brain regions composed the coupling term. Together with the intra-regional coupling by the IPSP signal from the inhibitory population, the total PSP input to the excitatory population is converted by a nonlinear sigmoid function *σ*_*e*_(*v*) given in Equation 8 below to an averaged firing density. The inhibitory population in region *n* received an input EPSP signal weighted by parameter *C*_*ie*_ from the excitatory population of the same region only, which was again converted to an averaged firing density by the following sigmoid function *σ*_*i*_(*v*):

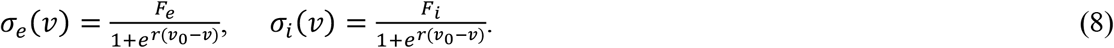

In Equations 8 of the mentioned sigmoid functions, the parameter *r* is a slope, *v*_0_ is a half of the maximal neural activity, and parameters *F*_*e*_ and *F*_*i*_ are the maximal firing densities of the excitatory and inhibitory populations, respectively. Parameter values of the considered two-population model (1)-(8) are given in Table 2.

**Table 2.** Parameter values of the electrical model and the Balloon-Windkessel model.

| Electrical model | Variables | Values | Balloon-Windkessel model | Variables | Values |
| --- | --- | --- | --- | --- | --- |
| Max. sigmoid (excitatory) | $F_e$ | 100 s<sup>-1</sup> | Echo time | $TE$ | 30 ms |
| Reciprocal of the time constant of the EPSP kernel | $a$ | 100<sup>a</sup> s<sup>-1</sup> | Mean-transit-time | $t_{MTT}$ | 2<sup>b</sup> s |
| Max. EPSP | $A$ | 3.25<sup>a</sup> mV | Net oxygen extraction fraction at rest | $E_0$ | 0.4<sup>b</sup> |
| Max. sigmoid (inhibitory) | $F_i$ | 50 s<sup>-1</sup> | Venous blood volume fraction | $V_0$ | 4<sup>b</sup> % |
| Reciprocal of the time constant of the IPSP kernel | $b$ | 50<sup>a</sup> s<sup>-1</sup> | Frequency offset for 3 T | $\vartheta_0$ | 80.6<sup>b</sup> s<sup>-1</sup> |
| Max. IPSP | $B$ | 22<sup>a</sup> mV | Ratio of intra/extra-vascular signal | $\varepsilon$ | 0.3<sup>b</sup> |
| Slope of sigmoid | $r$ | 0.56<sup>a</sup> mV<sup>-1</sup> | Sensitivity (regression slope) | $r_0$ | 25<sup>b</sup> s<sup>-1</sup> |
| 50 % neural activity | $v_0$ | 6<sup>a</sup> mV | Steady-state flow-volume relationship | $\alpha$ | 0.38<sup>b</sup> |
| Intra-regional coupling (from excitatory to inhibitory) | $C_{ie}$ | 6 | Rate constant for damped oscillations | $\kappa$ | 0.64<sup>b</sup> Hz |
| Intra-regional coupling (from inhibitory to excitatory) | $C_{ei}$ | 6 | Rate constant for damped oscillations | $\gamma$ | 0.32<sup>b</sup> Hz |
| Scaling factor | $R$ | 2.2 | Range of random initial conditions | $[s, f, v, q]$ | $[0, 1, 1, 1]$<sup>c</sup> |
| Amplitude of noise |  | 1.5 V/s<sup>2</sup> |  |  |  |

<sup>a</sup>Values from Jansen and Rit study.
<sup>b</sup>Values from Havlicek et al.
<sup>c</sup>Values empirically determined based on the trajectories generated by the Balloon-Windkessel model.
Abbreviations: IPSP = inhibitory postsynaptic potential; EPSP = excitatory postsynaptic potential

#### Simulated BOLD signals

We calculated the regional BOLD signals using the corresponding EPSP signals simulated by the electrical model (1)-(8) introduced in the previous section. Several examples of the time courses of the EPSP signals generated by the considered model and their power spectra are illustrated in Supplementary Fig. 1. Neurovascular coupling and hemodynamic responses constitute the process reflected in the Balloon-Windkessel (BW) model that was utilized to convert the simulated neural activity to BOLD signals,^58-60^ see details in the Supplementary material.

### Model validation: Neuroimaging and behavioral model fitting

In this study, we considered two model fitting approaches: neuroimaging model fitting and behavioral model fitting. The former is well known in the literature and consists of validation of the model via comparing simulated data against neuroimaging empirical data. In this study, the Pearson’s correlation coefficient between eFC and sFC (comparing the upper triangle without self-connections of the connectivity matrices) was calculated and denoted as goodness-of-fit (GoF) values. Searching for the maximal GoF in a given parameter space is a well-established approach for model validation in whole-brain modeling studies.^13-15^ In this study we optimized the coupling and delay model parameters to maximize the GoF value on a parameter grid of 64 × 43 points (64 global couplings and 43 global delays) densely covering the parameter plane, respectively. In addition, we also considered the connectivity relationship between eSC and sFC as for separate neuroimaging model fitting. In consequence, two types of neuroimaging model fitting (eFC versus sFC and eSC versus sFC) were used in this study. As this procedure fits the model to the connectivity derived from the empirical neuroimaging data, we termed it *neuroimaging model fitting*.

We also introduce *behavioral model fitting* as a procedure to validate a model against behavioral data, for example, optimizing the model to reflect some behavioral (phenotypical) properties to a best possible extent. In this study, we optimized parameters of the model to maximally differentiate between PD patients and HC subjects. For this, we calculated the effect size based on *z*-statistics of the Wilcoxon rank-sum two-tail test as given by the Rosenthal formula, i.e., the normal *z*-statistics divided by the square root of the number of observations^61^ of the difference between (neuroimaging) GoF values of the HC and PD subject groups. The effect size was calculated for every parameter point in the considered parameter space of 64 × 43 grid and represented as a parameter map. In this way we obtained a parameter landscape of the group differences and were able to investigate the differentiation of GoF values of PD patients from those of HC subjects. This parameter landscape reflects the relation of the model GoF to the behavioral data (in this study, to the differentiation based on clinical measures), and we thus used this approach as behavioral model fitting. To evaluate the parameter areas of significant group difference, we performed the Wilcoxon rank-sum two-tail test and obtained a corresponding *p*-value parameter map. Due to the multiple comparison over the parameter points, we applied the random-field thresholding scheme^62,63^ using a 2-dimensional Gaussian kernel smoothing. Subsequently, we obtained a z-score map and thresholded it to retain statistically significant parameter areas (alpha = 0.05). Finally, we searched for the optimal model parameters within the significant parameter areas corresponding to the maximal effect size. We considered two connectivity relationships (eFC vs. sFC and eSC vs. sFC) for the behavioral model fitting.

### Random sampling for optimal parameters

We performed a random sampling to test stability of the optimal parameter points for the behavioral model fitting. To do this, the stability of the results was assessed by a sex-balanced stratified subsampling. After a random sampling of 72 subjects (36 HC subjects and 36 PD patients) out of 116 subjects, we applied the behavioral model fitting on the sampled subjects and found optimal parameters corresponding to the largest effect size. The sub-sampling and the corresponding calculations were repeated 1000 times.

### Regularized (LASSO) logistic regression

The current task is to train a binary classifier (PD versus HC) using ten features (five connectivity relationships from two parcellation schemes), which is less dimension than observations (116 subjects). Thus, a logistic regression is applicable to the current study. To this end, we used a regularized logistic regression with the least absolute shrinkage and selection operator (LASSO) for training and classification of HC versus PD subjects.^64^ To avoid an overfit, the training error included the deviance and an L1-penalty.^65^ We used the *lassoglm* function for the logistic LASSO regression and the *glmval* function for predicted probability calculation in the Statistics and Machine Learning Toolbox of MATLAB R2020b.

### Confound regression for age-controlled features

We used a cross-validation (CV) scheme to train the logistic LASSO regression for PD classification. As for a degenerative disease,^66,67^ features for PD classification should be controlled by an age effect via confound regression. Due to a random sampling from the same cohort and the usage of the same data for the model validation and model training, it is important to prevent possible data leakage during the CV procedure, especially for behavioral model fitting as it uses across subjects. Otherwise, the trained models might be biased due to the usage of the results of the behavioral model fitting derived from PD classification against HC. In this respect, we followed the ideas of the cross-validated confound regression^68^ as illustrated in Fig. 2. Specifically, we applied the CV-consistent approach to features derived from the empirical result, neuroimaging, and behavioral model fitting. Accordingly, the subjects were split into training and test sets (Fig. 2, green and orange blocks in the outer loop), and the optimal parameter point of the behavioral model fitting was calculated on the training set at every iteration of the outer CV loop (Fig. 2, the green box with the circle 1). Then the respective connectome relationships were calculated for every subject. Next, the age was regressed for these subjects out (cross-validated confound regressions in Fig. 2, circles 1 and 2) from the obtained features of connectivity relationships used for subject classification. The optimal model parameters and the regression coefficients obtained for the training set were then used for the connectome calculation and the age regression for the test subjects.

**Figure 2.**
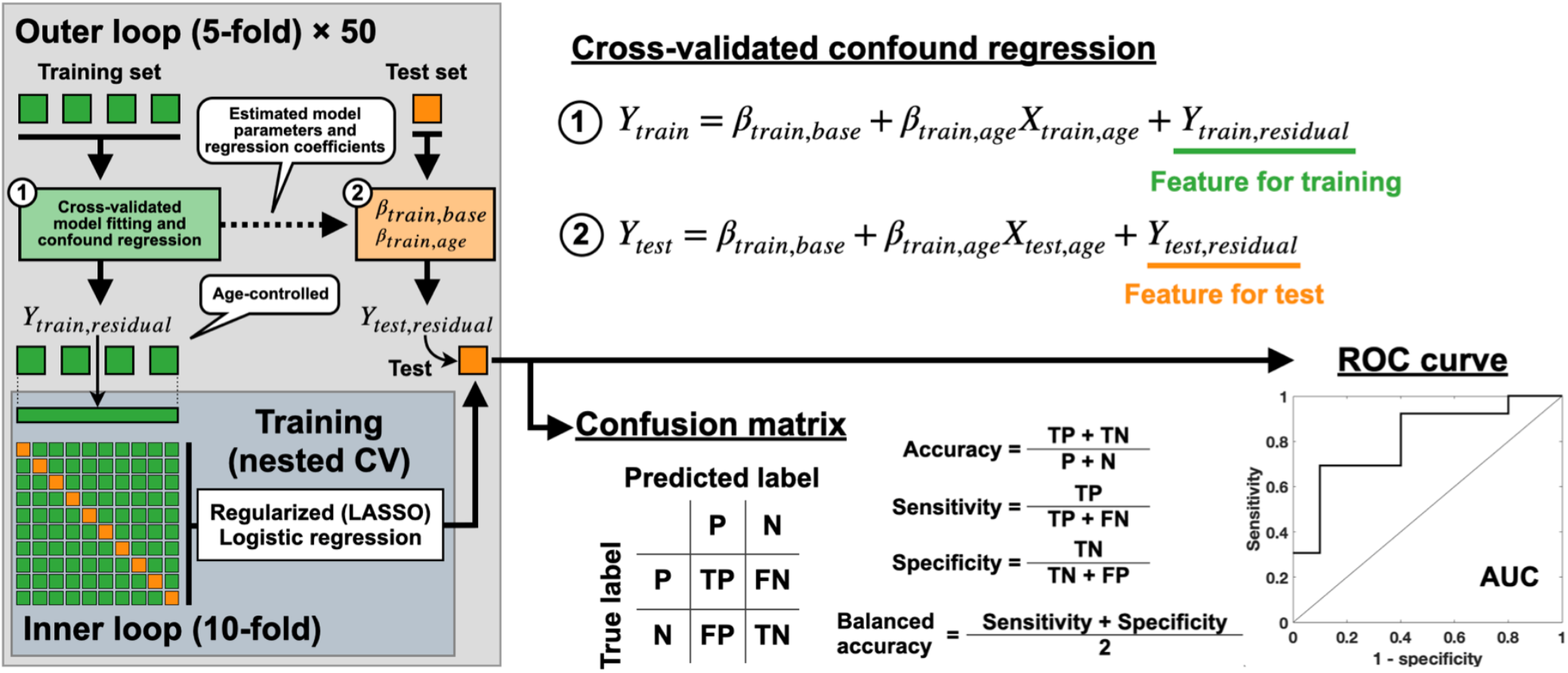
Schematic illustration of cross-validated model fitting, cross-validated confound regression and nested cross-validation (CV). The green boxes in the leftmost plot illustrate randomly split subject subgroups used for training of the model in the 5-fold outer loop and in the 10-fold inner loop. The orange box in the outer loop depicts the testing subject subgroup used for evaluation of the prediction performance of the trained model as given by accuracy, sensitivity, specificity, balanced accuracy and area under ROC curve. Abbreviations P: positive as patients, N: negative as controls, TP: true positive, FP: false positive, TN: true negative, FN: false negative, LASSO: least absolute shrinkage and selection operator, ROC: receiver operating characteristic, and AUC: area-under-curve.

### Nested cross-validation

In order to avoid over-optimistic results of CV,^69^ we used nested CV to train the logistic LASSO regression for PD classification (Fig. 2). In an outer loop, we randomly split the subjects into five subsets. One subset of 20% of subjects was considered as a test set (unseen subjects, the orange box in the outer loop in Fig. 2), and the other four subsets were pulled together and composed a training set (the green boxes in the outer loop in Fig. 2). As explained above, we first applied the cross-validated model fitting and confound regression to the features in the training set (Fig. 2, the green box with the circle 1). Subsequently, the training set (age-controlled) was split into ten subsets for the nested CV in the inner loop. A logistic LASSO regression model was trained with the hyperparameters minimizing the 10-fold CV error. This model was then applied to predict the test set. As follows from the aforementioned, the age-controlled training and test sets were used for model training and prediction, respectively. The training and test procedure we performed can be summarized as follows:

1. Randomly split the entire subject cohort into five subgroups.
2. Select one group as a test set and compound the others into a training set.
3. Perform the cross-validated (behavioral) model fitting using the training set and extract respective connectome relationships corresponding to the optimal model parameters.
4. Perform the cross-validated confound (age) regression for the training set from the features based on the connectome relationships used for classification.
5. Train the logistic LASSO regression model in the inner loop with 10-fold CV that minimizes errors of the prediction model.
6. Apply the trained best model to predict the test set with age regressed, where the optimal model parameters of the model fitting and age regression coefficients obtained for the training set were used (Fig. 2, the dashed arrow in the outer loop).
7. Calculate the model performance using a confusion matrix and an ROC curve.
8. Perform steps 2-7 for the other four subsets split in step 1 as an outer CV loop (5 prediction results).
9. Repeat steps 1-8, 50 times (250 prediction results in total).

### Evaluation of prediction performance

For PD classification based on the discussed machine learning approach we considered five features for each of the two parcellation schemes (Schaefer and Desikan-Killiany atlases), i.e., 10 features in total: corr(eFC, eSC) as an empirical feature, corr(eFC, sFC) and corr(eSC, sFC) as simulated features for each model fitting, i.e., the neuroimaging model fitting and the behavioral model fitting. To investigate the impact of simulated results on the PD prediction, we composed the considered features into three conditions: *(i)* empirical features only (shuffle simulated features), *(ii)* simulated features only (shuffle empirical features), and *(iii)* all features (no shuffling). The shuffling was performed by a random re-distribution of the values of a given feature among subjects such that the correspondence of the feature to individual subjects was destroyed. By focusing on some features (connectome relationships and parcellations), the other features were shuffled. For example, to focus on the empirical features for the Schaefer atlas, four simulated features (eFC vs. sFC and eSC vs. sFC for two model fitting modalities) of the Schaefer atlas and all five features (one empirical and four simulated features) of the Desikan-Killiany atlas were shuffled. The shuffling was performed for every feature separately randomizing feature values across subjects while retaining distributional properties (see Supplementary Fig. 2 for details). After feature selection, model training and application of the trained model to the unseen test subject set, we calculated a confusion matrix from the prediction results and plotted a receiver operating characteristic (ROC) curve.^70^ The latter was calculated from the prediction results obtained by varying a subject classification threshold of a predicted probability from 0 to 1. Then, we calculated the prediction performance (accuracy, sensitivity, specificity, and balanced accuracy) and the area-under-curve (AUC) of the ROC curve.

In addition to the prediction considering the cross-validated confound regression with subjects’ ages using the entire cohort, we also applied the same approach to a balanced subject configuration by excluding 17 oldest PD patients from 116 subjects. Thus, the balanced cohort has no significant age difference between PD and HC groups in balanced group sizes (see Supplementary Table 1). Subsequently, we analyzed the prediction performances from the balanced subject cohort (99 subjects).

We also scrutinized the prediction probabilities for individual subjects to evaluate the model performance. Here, the trained model estimated the predicted probabilities for each subject in the test set. Subsequently, we calculated a fraction of actual positives and showed relationships using probability calibration. The ideal case is to have the same values of the fraction of positives and the predicted probability, i.e., the graph should align to the diagonal. In clinical applications, the tight correspondence between predicted probabilities and the fraction of actual positives provide high trustworthiness for diagnosis.^71^ To this end, we used Brier score^72^ to calculate the mean-squared error of each predicted probability against an ideal case. We also used Wasserstein distance to show how much cost is required to turn a given distribution of the predicted probabilities into a uniform one.^73^ In other words, this metric was used to evaluate how well predicted probabilities were uniformly distributed. Thus, a lower Wasserstein distance means that the predicted probabilities are relatively better calibrated than that of a higher one. Accordingly, we further evaluated the model performance regarding individual predicted probabilities in addition to the integrative performance from the confusion matrix.

### Data availability

The raw empirical data used in this study are not immediately available for public sharing because the given informed consent of included patients did not include public sharing. The simulated data that support the findings of this study are available from the corresponding author upon a reasonable request.

## Results

In this study, we investigated the application of simulation results of whole-brain dynamical models to PD classification using relationships between empirical and simulated connectomes as features. The whole-brain dynamical model of the Jansen-Rit type was used to simulate the electrical neuronal activity and was validated against empirical data by means of neuroimaging or behavioral model fitting. Accordingly, we calculated the connectome relationships involving the simulated connectomes corresponding to the optimal model parameters of the two fitting modalities and used them as features for PD classification. We show that complementing the empirical data by simulated data improves the prediction performance as compared to the case where only empirical data were used.

### Neuroimaging model fitting

We calculated sFC using simulated BOLD signals for each parameter point and obtained the similarity (Pearson’s correlation) values between eFC and sFC. Fig. 3 shows the corresponding values in the delay-coupling (*τ*_*global*_, *C*)-parameter space averaged over all the subjects, which defines an average GoF parameter landscape (Fig. 3 A and E). We calculated eFC and sFC for the different frequency ranges of the corresponding filtered BOLD signals, i.e., NF, BF, LF, and HF conditions (see Methods for details). The profiles of the parameter landscapes were different between the considered brain atlases. The Schaefer atlas showed a unimodal distribution containing maximal GoF values (the dashed circle in Fig. 3 A) for the optimal global delays in the biologically feasible range^74^ from 0.06 to 0.25 s/m (Fig. 3 D). On the other hand, the maximal GoF for the Desikan-Killiany atlas posited a bi-modal distribution (the dashed circles in Fig. 3 E) with well separated peaks along the global coupling parameter (Fig. 3 G). Moreover, stronger global coupling of the maximal GoF values was accompanied by a widespread of global delays (the upper dashed circle in Fig. 3 E) that may get out of the biologically feasible range as compared to the weaker global couplings (the lower dashed circle in Fig. 3 E).

**Figure 3.**
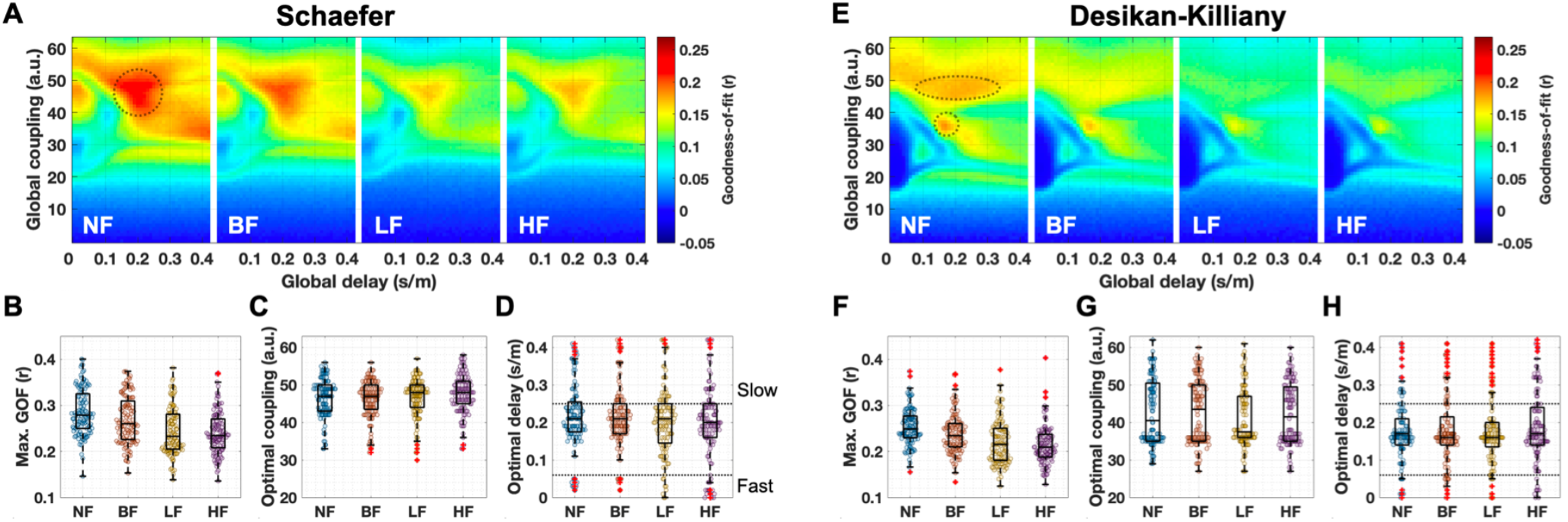
Results of the neuroimaging model fitting. (A-D) The Schaefer atlas and (E-H) the Desikan-Killiany atlas. (A, E) Parameter landscapes of the similarity (Pearson’s correlation) between empirical FC and simulated FC, i.e., goodness-of-fit (GoF) values averaged over the entire subject cohort. The landscapes are illustrated for each filtering condition (NF, BF, LF, and HF, see Methods for details). The dashed circles delineate the hills with large GoF values. Distributions of (B, F) the maximal GoF values, (C, G) optimal coupling parameters and (D, H) the respective optimal delays corresponding to the maximal GoF values for each filtering condition. The dashed horizontal lines in plots (D, H) indicate the biologically feasible delay range regarding the electrophysiological conduction speed. The middle lines in interquartile box plots indicate medians of distributions, and the red pluses are the outliers. Abbreviations NF: no filtering, BF: broad band ([0.01,0.1] Hz), LF: low-frequency band ([0.01,0.05] Hz, HF: high-frequency band ([0.05,0.1] Hz).

Furthermore, we observed that applying temporal filtering to BOLD signals diminishes GoF values over the entire parameter landscape. In particular, the narrow frequency bands (LF and HF) resulted in significantly lower maximal GoF values than for the cases of the broader (BF) or the entire frequency (NF) range, see Table 3 for statistical results.

**Table 3.** **Comparisons between goodness-of-fit values of the considered filtering conditions (Bonferroni corrected *p*-values of the Wilcoxon signed-rank two-tail test) and the corresponding effect sizes by Rosenthal formula. Bold fonts indicate that the goodness-of-fit values are significantly different between filtering conditions.**

| **$p$ (effect size)** | **NF vs. BF** | **NF vs. LF** | **NF vs. HF** | **BF vs. LF** | **BF vs. HF** | **LF vs. HF** |
| --- | --- | --- | --- | --- | --- | --- |
| Schaefer | **0.000 (0.70)** | **0.000 (0.84)** | **0.000 (0.86)** | **0.000 (0.81)** | **0.000 (0.70)** | 0.998 (0.04) |
| Desikan-Killiany | **0.000 (0.66)** | **0.000 (0.77)** | **0.000 (0.85)** | **0.000 (0.69)** | **0.000 (0.70)** | 0.838 (0.10) |

Abbreviations: NF = no filtering; BF = broad band ([0.01,0.1] Hz); LF = low-frequency band ([0.01,0.05] Hz); HF = high-frequency band ([0.05,0.1] Hz).

### Effect size of group comparisons for behavioral model fitting

The behavioral model fitting resulted in effect sizes of group difference between HC and PD (Fig. 4 A-B for eFC-sFC correlation, see Supplementary Fig. 3 for eSC-sFC correlation). Furthermore, we also observed that the distributions of the optimal parameter points corresponding to the maximal effect sizes to be densely concentrated in the parameter space across repeated sub-sampling (1000 times) and filtering conditions (Fig. 4 C-D, distributions in blue). Interestingly, the distributions of the optimal parameters derived from the behavioral model fitting were strikingly different from those determined by the neuroimaging model fitting (Fig. 4 C-D, distributions in orange for the neuroimaging and blue for the behavioral). Both sets of the optimal parameters are located in the biologically plausible range of time delay.^74^

**Figure 4.**
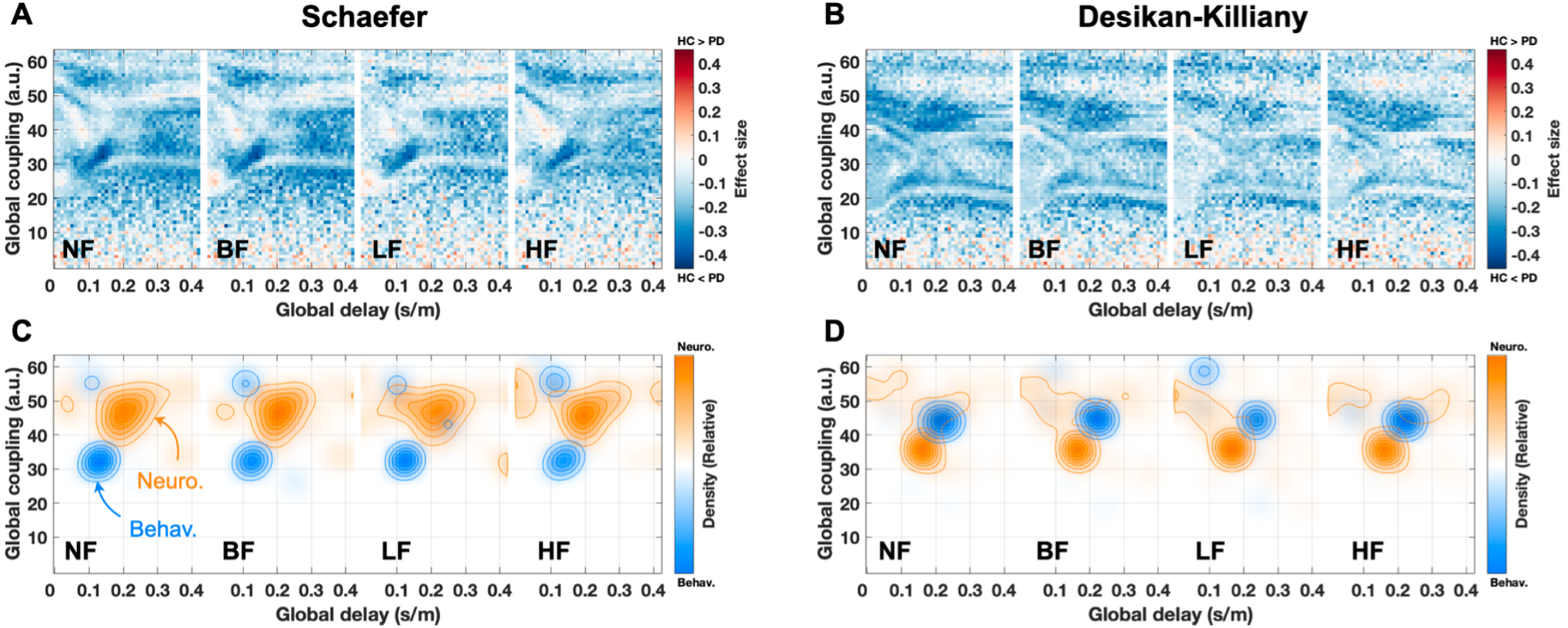
Parameter maps of the effect size of the difference between goodness-of-fit (GoF) values (eFC-sFC correlation) of healthy and Parkinsonian groups used for the behavioral model fitting. The filtering conditions are indicated in the plots for (A) the Schaefer atlas and (B) the Desikan-Killiany atlas. Effect sizes in the (*τ*_*global*_,*C*)-parameter plane were calculated by a non-parametric Wilcoxon rank-sum two-tailed test between patients and controls in the GoF values for each parameter point. (C, D) Distributions of optimal parameters derived from the neuroimaging model fitting (orange, all subjects, n=116) and the behavioral model fitting (blue, repeated sub-sampling, n=1000) for (C) the Schaefer atlas and (D) the Desikan-Killiany atlas. Abbreviations PD: Parkinson’s disease, HC: healthy controls, NF: no filtering, BF: broad band ([0.01,0.1] Hz), LF: low-frequency band ([0.01,0.05] Hz, HF: high-frequency band ([0.05,0.1] Hz).

### Group difference between healthy controls and patients

The empirical structure-function relationships corr(eFC, eSC) for HC and PD subject groups were found to be from distributions with different medians for the Schaefer atlas and all considered filtering conditions and for the LF condition only for the Desikan-Killiany atlas (Fig. 5, the first row). The group differences obtained by involving the simulated connectomes of the neuroimaging model fitting were small and non-significant for both atlases and all filtering conditions (Fig. 5, the second and third rows). On the other hand, for behavioral model fitting we observed that PD patients exhibited stronger agreements between empirical and simulated connectomes than HC subjects and can thus be better differentiated from HC (Fig. 5, the fourth and fifth rows).

**Figure 5.**
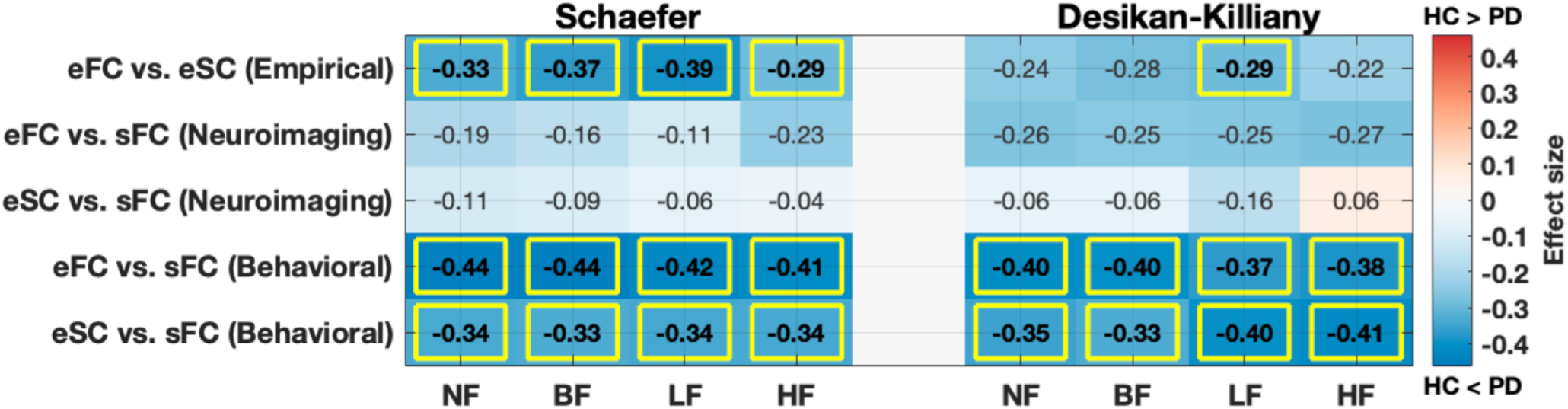
Differentiation between healthy and Parkinsonian subjects as reflected by the relationships between empirical and simulated connectomes. (Left) The Schaefer atlas and (Right) the Desikan-Killiany atlases. The simulated connectomes are calculated for the optimal model parameters of the neuroimaging and behavioral model fitting as indicated on the vertical axis. Summary tables of the effect sizes (numbers) of the differences between Parkinsonian and healthy subject groups are calculated by the Rosenthal formula and shown in blue (negative) for HC > PD and red (positive) for HC < PD. The significant cases are indicated by yellow rectangles as given by the Bonferroni corrected *p*-values of the Wilcoxon rank-sum two-tail test. Abbreviations PD: Parkinson’s disease, HC: healthy controls, NF: no filtering, BF: broad band ([0.01,0.1] Hz), LF: low-frequency band ([0.01,0.05] Hz, HF: high-frequency band ([0.05,0.1] Hz).

Temporal filtering may influence the group differences for the empirical and also for the simulated connectomes as illustrated in Fig. 5, see the first row for the Desikan-Killiany atlas, in particular, and Supplementary Fig. 4. In addition, we calculated the explained variances of the five connectivity relationships between each other for the same and different filtering condition, which resulted in relatively low similarities for the simulated results (Supplementary Fig. 5). Accordingly, the temporal filtering can influence the considered connectivity relationships and may lead to dissimilar patterns of connectome relationships across subjects.

### Prediction performance

We used the five whole-brain connectivity relationships as features for PD classification using machine learning based on the logistic LASSO regression algorithm. The feature space constituted three feature conditions with ten features (five connectivity relationships for two atlases), see Supplementary Fig. 2. After the nested CV, the trained best models were relatively well balanced with a slight tendency towards overfitting for some of the used performance measures (13.4% decreased balanced accuracy and 1.1 % decreased AUC of test performance from training one, see Supplementary Fig. 6).

Fig. 6 shows the prediction performance for each of the investigated conditional cases of brain parcellations, frequency bands and feature conditions. The first important observation is that an involvement of the simulated connectomes can improve the classification of PD and HC, see Fig. 6 and compare blue bars (empirical features) to red bars (simulated features) and to yellow bars (all features) (see Supplementary Fig. 7 for the differences). In the latter case, where the empirical features are complemented by the simulated ones, the prediction performance can only be enhanced as compared to purely empirical features, which we observed for most feature conditions and performance measures (Fig. 6). Interestingly, the performance further improved when using features from both atlases (Fig. 6 and Supplementary Fig. 7).

**Figure 6.**
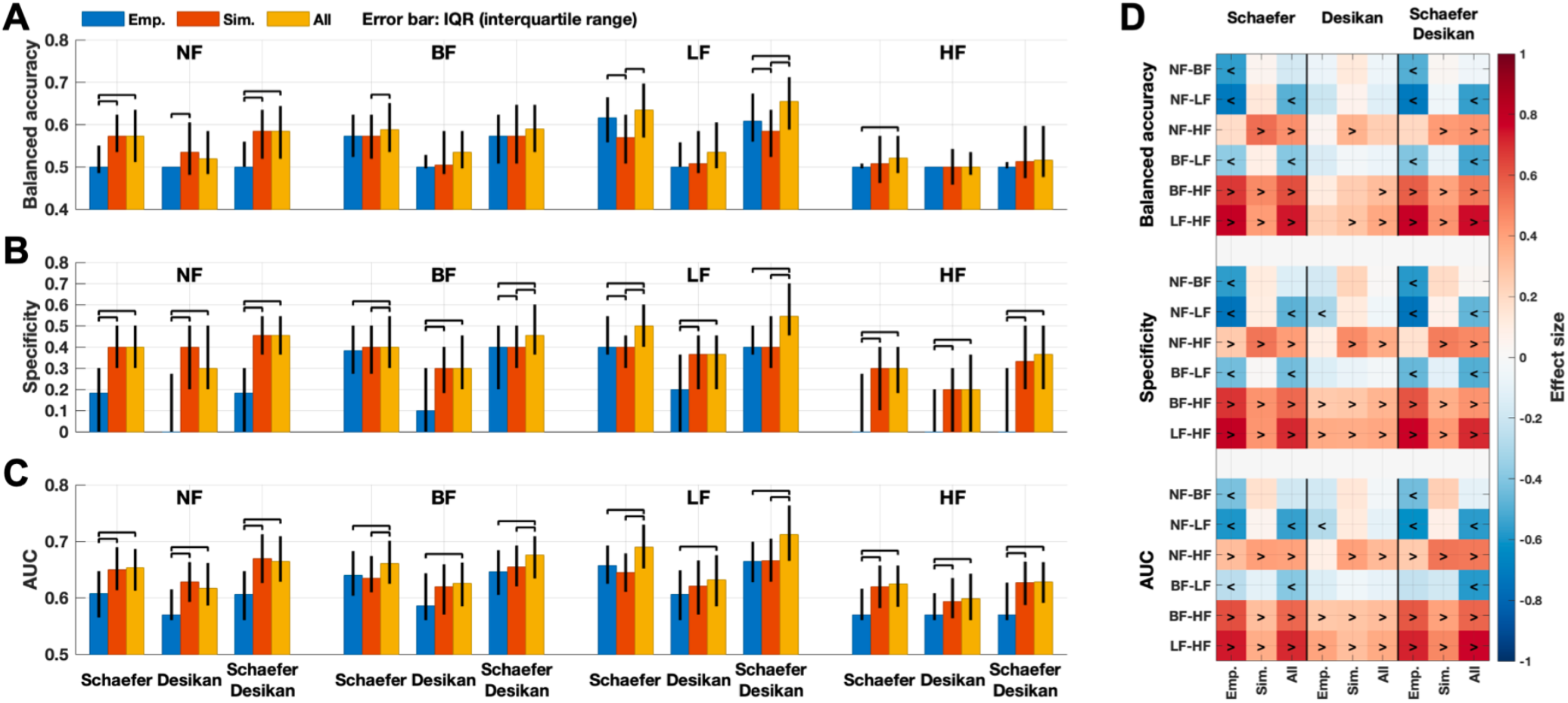
Summary of the performance of Parkinson’s disease classification using the three different feature conditions: empirical features (blue bars), simulated features (red bars), and all features (yellow bars) after incorporating the age controlling and the behavioral model fitting during the nested cross-validation (Fig. 2). Median values of the balanced accuracy, specificity and area-under-curve (AUC) of the receiver operating characteristics (ROC) curves for all considered parcellations and filtering conditions are shown in each panel for (A) balanced accuracy and (B) specificity and (C) AUC. The error bars indicate the interquartile range across iterations of the outer loop of the nested cross-validation procedure (Fig. 2). The black lines connecting two conditions indicate significantly different performance between feature conditions. (D) Effect sizes between filtering conditions for each feature condition. The signs ‘<’ and ‘>’ indicate which condition is significantly larger than the other. For example, ‘<’ sign for ‘NF-LF’ indicated on the vertical axes means NF < LF for a given performance indicated on the horizontal axes. The Wilcoxon signed-rank two-tail test was used for comparisons across feature and filtering conditions (Bonferroni corrected statistics). The Desikan-Killiany atlas is shorten as “Desikan”. Abbreviations NF: no filtering, BF: broad band ([0.01,0.1] Hz), LF: low-frequency band ([0.01,0.05] Hz, HF: high-frequency band ([0.05,0.1] Hz).

We also investigated how the prediction performance varies depending on the filtering conditions (Fig. 6 D). The effect of the temporal filtering was prominent for the empirical features for the Schaefer atlas, where the performance was significantly increased for the LF condition compared to the others (Fig. 6 D, the Emp. column for the Schaefer atlas). On the other hand, the HF condition showed low performances on the empirical features, in particular, with very low specificities down to 0 (Fig. 6 B and D) and very high sensitivities up to 1 (Supplementary Fig. 8), where the LF filtering seems again to be a beneficial condition for PD prediction. Summarizing, the temporal filtering conditions influenced the model performance, and the LF band-pass filtering resulted in the most effective prediction relying on the connectome relationships. The other considered narrow-band HF filtering condition is not advisable for PD classification. However, involving the simulated connectomes is still of advantage also under this condition as compared to using only empirical features.

We also compared the prediction performance when the simulated connectomes obtained from the neuroimaging and behavioral model fitting were considered separately. This resulted in two additional feature conditions (see Supplementary Fig. 8). The neuroimaging model fitting in most cases led to a weaker prediction performance compared to the behavioral model fitting or to the composite case when the features of both fittings are merged. This justifies the introduction of the behavioral model fitting for subject classification.

Furthermore, we applied the current approach to the balanced subject configuration (99 subjects, see Supplementary Table 1 for the demography). The prediction performance was consistent with the main findings of the entire cohort (116 subjects, Fig. 6). In other words, complementing empirical data with simulated results using the LF filtering involving the multi-parcellation (concatenating both atlases) is advisable for PD classification (Supplementary Fig. 9).

Fig. 6 shows the well-known measures characterizing the prediction performance as median values and interquartile ranges of distributions. Although these measures clearly reflect how well the machine learning approach is commonly working, we may also be interested in how every test is performing for classification of individual unseen subjects. In this respect, Fig. 7 illustrates the results of classification/prediction probabilities of all tests performed on individual subjects from the test sets. The prediction probabilities were collected and related to the probability calibration curves.

**Figure 7.**
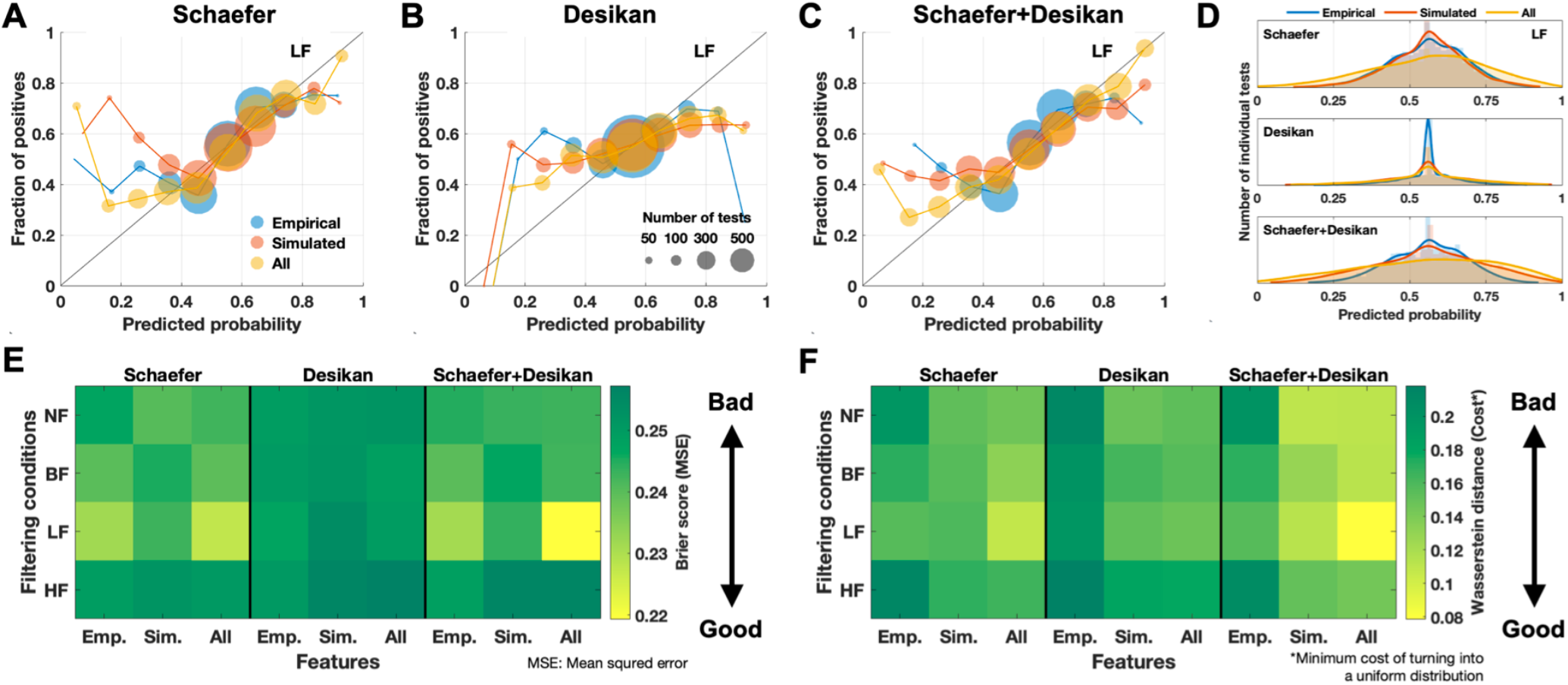
The performance of the trained prediction model regarding the predicted probabilities for individual subjects. The case of the low-frequency bandpass filtering condition (LF) is illustrated. (Top row) Plots of the probability calibrations for (A) the Schaefer atlas, (B) Desikan-Killiany atlas, and (C) multiple atlases, where the fraction of true positives is plotted versus the probability of them predicted by trained model for individual subjects. The sizes of circles indicate the number of individual subject tests for the three considered feature conditions as indicated in the legend in plot (A). (D) Histograms of the predicted probabilities of the low-frequency filtering condition for each feature condition as indicated in the legend. (E) Table of the Brier scores (mean-squared error) for all considered filtering and feature conditions. (F) Tables of the Wasserstein distances between distributions of predicted probabilities and a uniform distribution for all conditions. Abbreviations Desikan: Desikan-Killiany, Emp.: empirical features, Sim.: simulated features, and All: empirical and simulated features, NF: no filtering, BF: broad band ([0.01,0.1] Hz), LF: low-frequency band ([0.01,0.05] Hz, HF: high-frequency band ([0.05,0.1] Hz).

We can interpret the probability calibration plots (Fig. 7 A-C) according to two aspects. Feature conditions using simulated results (red and yellow curves) resulted in predictions that are more closely aligned to the ideal case (the diagonal black line) than the empirical relationship. Indeed, for the Schaefer atlas and the multi-parcellation case the distance to the diagonal as given by the mean-squared error of the predicted probabilities against the actual classes calculated according to the Brier score^72^ is minimal for the composed features including the empirical and simulated connectomes for the LF filtering condition (Fig. 7 E). As a second aspect, the prediction probabilities derived from the empirical features are more narrowly distributed around 0.5 (blue curves in Fig. 7 D) compared to the case of all features (yellow curves in Fig. 7 D). This can be quantified by the minimum cost of turning the observed distribution into a uniform distribution using the Wasserstein distance^73^ (Fig 7 F). In the latter case, the predicted probabilities derived from all features show widely spreading distributions also reaching the low and high probability values, which indicate a high confidence.^71^ In other words, in our predictive modeling, the prediction results where the empirical data are complemented by simulated features were better calibrated in some cases as compared to the case of the empirical data only (Fig. 7 C). As mentioned above, the Wasserstein distance in Fig. 7 F clearly shows which filtering condition and which feature condition can be the best beneficial configuration for PD classification. In particular, the LF filtering of the BOLD signals and involving of the simulated connectomes together with the empirical ones for the Schaefer atlas and multi-parcellation case can improve the prediction results and the confidence of the prediction model. The same conclusion was drawn above based on the Brier scores, which confirms its robustness and which may be of relevance for application of the discussed modeling and prediction approaches to clinical data and disease diagnosis.

## Discussion

The main objective of this study is to effectively apply whole-brain dynamical modeling and the derived simulated connectomes to PD classification. Whole-brain simulations allow us to explore various regimes of brain dynamics corresponding to different values of free model parameters. To extract features from the simulated results, it is essential to evaluate which model fitting is appropriate. The detected optimal model parameters can differ when we use different model-fitting approaches. In other words, whole-brain dynamics with proper model parameters can disclose group differences between PD and HC subjects and provide a way to extract effective features for PD classification. In this study, we introduced the behavioral model-fitting approach and showed that it captures differences between PD and HC better than the conventionally used neuroimaging model fitting approach. Then we applied it to PD classification. Based on our findings, we can conclude that using a proper model validation in the whole-brain dynamical modeling may provide effective features to machine learning, and it provides information complementary to empirical features.

In addition to the whole-brain dynamical modeling for classification, data processing is also important because as we have shown different data processing influences model validation.^6,30,35,75^ In this respect, we investigated how temporal filtering of BOLD signals and brain parcellation influence empirical and simulated results regarding the model fitting, group difference and prediction performance. Based on our results, we can conclude that the resting-state whole-brain simulations with an appropriate data processing and model validation reflect personal traits of individual subjects better, which may contribute to disease classification based on the whole-brain connectivity relationships with potential relevance in medicine.

### Effect of temporal filtering on model fitting and prediction

The effect of temporal filtering on functional MRI has been in the focus of neuroimaging research for a long time.^76-79^ One related study considered different temporal filters for MRI data processing and reported distinguishable BOLD dynamics in task-driven and resting-state brain activity between low and high frequency bandpass filtering.^38^ Furthermore, temporal filtering can influence the classification performance for patients with Alzheimer’s disease as compared across several low- and high-band pass filtering conditions.^39^ In this study we found that the neuroimaging model fitting resulted in significantly different distributions of the maximal GoF values for individual subjects for different filtering conditions. Furthermore, the empirical structure-function connectivity relationship and the maximal GoF values of the neuroimaging model fitting were diminishing for the narrower filtering bands (Supplementary Fig. 4).

Another study investigated PD classification via machine-learning on brain networks derived from the empirical resting-state FC with a high-pass temporal filtering (> 0.01 Hz) of BOLD signals,^36^ which corresponds to the case of the NF condition in our study. According to our prediction results, we suggest to consider the low-frequency bandpass filtering, i.e., the LF condition, which can improve the differentiation and classification of PD also for the case when only empirical features were used.

An appropriate selection of the filtering condition (broad- or narrow-, high- or low-frequency band) appears to be important for the prediction performance as reflected by several integrative measures considered in this study. In particular, a detailed evaluation of individual tests indicates that selecting a proper bandpass filter of the empirical and simulated BOLD signals can improve the prediction performance (Fig. 6 and Fig. 7). Thus, the involvement of simulated connectomes, especially, in combination with empirical ones, is a great advantage by means of the individual probabilities closer approach the ideal case as compared to the purely empirical feature space.

### Biophysical interpretation of model parameters

Under the assumption that the resting-state brain activity is governed by a complex dynamical system, we can interpret the optimal model parameters of the neuroimaging model fitting as parameters of that system with potential neuroscientific/physical meaning. Despite the optimal parameters determined by model validation, it can differ when a given model fitting approach changes observed in our previous studies^6,30^ and the results in the current study (Fig. 4 C-D). Furthermore, the parcellations also impact on the locations of the optimal parameters. For instance, the optimal global coupling parameters derived from the behavioral model fitting had weaker optimal couplings than those from the neuroimaging model fitting for the Schafer atlas (Fig. 4 C). On the other hand, the situation for the Desikan-Killiany atlas is opposite (Fig. 4 D).

In our model, we used the reconstructed path lengths of the tractography streamlines in the white matter, which approximate the actual lengths of the anatomical axonal connections in the brain. The considered model simulates the electrical activity of the excitatory and inhibitory neuronal populations in the brain regions as reflected by the dynamics of the respective PSP signals. We can thus evaluate and interpret the optimal model parameters of the propagation of the simulated electrical signals (EPSP) along the brain pathways. We, in particular, found that the neuroimaging model fitting resulted in the optimal delay of the signal propagation in the electrophysiologically plausible range^74^ (Fig. 3). This confirms the applicability of the used dynamical model for simulating brain dynamics. Furthermore, the optimal delay of the behavioral model fitting obtained at the repeated subsampling for different subject configurations is located in the same biologically reasonable range as well, which validates the behavioral model fit (Fig. 4 C-D). Further parameters of the considered model and the simulated electrical PSP signals (Table 2) may have biologically plausible interpretations and ranges. Here we may mention, for example, the excitation-inhibition balance of the intra-regional coupling or the time constants responsible for control of slow or fast oscillations of electrical neuronal activity.

In PD research, a neural model generating such oscillations in a certain frequency range is essential to engage the pathological neural activity during rest. Previous studies reported that the resting-state cortico-cortical FC of PD patients changed in 8-10 Hz range (in the alpha-rhythm) for early-stage and moderately advanced PD patients,^80^ and cortico-cortical coupling between 10 Hz and 35 Hz correlated with the severity of PD in the electroencephalogram study.^81^ High oscillatory synchrony in the basal ganglia at frequencies of 8-35 Hz was also associated with PD based on spectral power changes between off- and on-drug (levodopa dose).^82^ With this respect, we may also investigate the relationship between frequencies of neural activity and models varying scale factor *R* of the current whole-brain dynamical model.

### Exploring parameter landscapes

The neuroimaging model fitting is a well-established model validation as though maximizing GoF values of the model is the main objective of the model validation. Nevertheless, brain dynamics for non-optimal model parameters may also provide additional useful properties. They can contribute to the application of the dynamical models to analyze the brain and behavior. In particular, brain modeling with virtual brain or *in silico* models for brain abnormality has been used for clinical purposes.^26-28^ To this end, we explored the parameter landscapes of GoF values and searched for parameter points that provide optimal GoF values to effectively answer the current research question. As we reported in the results, there exist hotspots of the densely located optimal model parameters, where either neuroimaging or behavioral model fitting is the most effective, although these hotspots may not coincide (Fig. 4 C-D, the distributions in blue and orange). Therefore, systematical exploration of parameter landscapes allows us to find proper model parameters for a given purpose, which may be different in locations and other properties from one model application and research question to another. Accordingly, we conclude that exploring parameter landscapes of the whole-brain dynamical models using behavioral/phenotypical measures might reveal optimal model parameters best suited for research goals related to the inter-individual variability and prediction approaches.

### Classification of Parkinsonian patients

In this study, we did not aim at obtaining the highest prediction accuracy, which might have required an extensive testing of many simulation and prediction conditions, feature spaces, and learning algorithms. Nevertheless, the obtained prediction performance (65.2 % as median accuracy using empirical features) is comparable with that reported, for example, in the study of Pläschke *et al*.^36^ which had a median accuracy of 65.5 % over considered brain networks.

When we consider the simulated data for PD classification, the features from the neuroimaging model fitting had much lower performance in most considered cases as compared to the features from the behavioral model fitting (Supplementary Fig. 8). Therefore, we suggest that the behavioral model fitting can be used to validate the model against behavioral data for probing the simulated whole-brain dynamics to improve the model correspondence to phenotypical characteristics of subjects and prediction results. Such an approach may be of crucial importance in clinical research, and the reported results showed first promising confirmations.

In this study, we also explored the impact of a few data processing choices and model simulation on the differentiation and prediction performance. For example, composing predictive features including empirical and simulated connectomes from multiple brain atlases can provide complementary features leading to even better prediction performance (Supplementary Fig. 7). We further showed that also filtering conditions of empirical and simulated BOLD signals can play an important role in model validation and subject classification, where in particular, prediction specificity may vary significantly across filtering conditions as well as the number of false positives of the trained model can be reduced by appropriate filtering (Fig. 6).

The modern neuroimaging research dedicated to prediction analysis and based on the machine-learning techniques has shown an enhanced performance for clinical data and in radiology in particular.^83,84^ Those predictive results and developed approaches have faced the issue of translation of their analysis and interpretation of the obtained outcomes to clinical application.^85^ In this respect, the current study illustrated the characteristics of individual prediction probabilities to bridge a gap between modeling and prediction results and their translation for diagnosis in clinical research. The analysis included in the present study explored the calibration of the predicted probabilities for individual subjects and provided additional reliable information for interpretation of the classification results. This can be achieved when the prediction probabilities are considered at the level of individual subjects, for example, when new unseen patients should be tested for diagnostic purposes. Furthermore, the discussed probability analysis delivered additional evidence that the whole-brain simulation results can be useful for complementing empirical data for prediction and classification in clinical research. Consequently, involving the whole-brain dynamical models in the training of machine-learning models can improve individual prediction, which can potentially help a clinician to better gauge a diagnosis during the examination of individual patients.

### Future work

For further studies, other phenotypical properties can be used for the behavioral model fitting, for instance, age or sex. Of course, cognitive or clinical scores such as the Montreal cognitive assessment, Mattis dementia rating scales, unified Parkinson’s disease rating scales are also applicable. The suggested approach of the behavioral model fitting is similar to the brain mapping of various behavioral or phenotypical measures on the cortical surface and can thus be generalized. In other words, we can map the parameter space using cognitive or clinical scores, which can be referred to as *phenotypical mapping* on the model parameter space like the behavioral model fitting that we introduced in the present study.

### Summary

We simulated whole-brain resting-state dynamics and calculated the relationships between structural and functional empirical and simulated connectomes for a variety of conditions of data processing options including brain parcellation and temporal filtering of BOLD signals. We introduced the behavioral model fitting paradigm and found that the ensuing modeling results can lead to an enhanced differentiation of disease and control groups and improved classification of Parkinsonian patients by machine-learning approaches. We showed that band-pass filtering in the low-frequency band can have a beneficial effect on the prediction performance. On the other hand, the high-frequencies of the empirical and simulated BOLD signals should be considered with care and may not immediately be recommended for subject-level classification. In addition, we demonstrated that the prediction performance can differ where we use different or multiple brain parcellation schemes. Our findings can contribute to a better understanding of empirical and simulated whole-brain dynamics and its relationship to disease. They further suggest an avenue for application of the results of whole-brain simulations for cognitive or clinical investigation of inter-individual differences and disease diagnosis.

## Acknowledgements

The authors greatly acknowledge the contribution of Dr. Christian Mathys, Dr. Martin Südmeyer, and Dr. Christian Hartmann for the assessment of the Parkinson’s disease data. The authors are also grateful to Shraddha Jain for the MRI data quality check. The authors gratefully acknowledge the computing time granted through JARA on the supercomputer JURECA at Forschungszentrum Jülich.

## Authorship contribution statement (CRediT)

**Kyesam Jung:** Conceptualization, Data curation, Formal analysis, Investigation, Methodology, Software, Validation, Visualization, Writing – Original draft preparation, Writing – Review & editing. **Esther Florin:** Conceptualization, Supervision, Validation, Writing – Review & editing. **Kaustubh Patil:** Methodology, Validation, Writing – Review & editing. **Julian Capers:** Data curation. **Christian Rubbert:** Data curation. **Simon Eickhoff:** Conceptualization, Funding acquisition, Project administration, Resources, Writing – Review & editing. **Oleksandr Popovych:** Conceptualization, Funding acquisition, Methodology, Project administration, Resources, Software, Supervision, Validation, Writing – Original draft preparation, Writing – Review & editing.

## Abbreviations

AUC: area-under-curve
BF: broad-frequency band
BOLD: blood oxygenation level-dependent
BW: Balloon-Windkessel
CV: cross-validation
DWI: diffusion-weighted image
EPSP: excitatory postsynaptic potential
FC: functional connectivity
FOD: fiber oriented distribution
GoF: goodness-of-fit
HC: healthy controls
HF: high-frequency band
IPSP: inhibitory postsynaptic potential
LASSO: least absolute shrinkage and selection operator
LF: low-frequency band
NF: no filtering
PD: Parkinson’s disease
PL: path length
PSP: postsynaptic potential
ROC: receiver operating characteristic
SC: streamline count
T1w: T1-weighted image
WBT: whole-brain tractography

## Supplementary material

### Simulated BOLD signals

The neurovascular coupling describes that the changes of the induced signals *s*(*t*) driven by the EPSP input link to the changes in the cerebral blood flows (CBF) *f*(*t*) as the blood inflow,

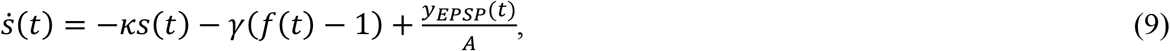

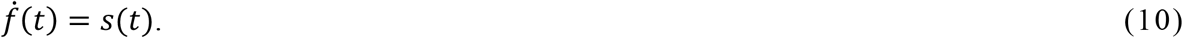

Equations 9 and 10 govern the dynamics of the induced signal and CBF, respectively. Parameters *κ* and *γ* are the rate constants that regulate ultra-slow endogenous fluctuations at around 0.09 Hz.^1^ The normalized neural response, i.e., *y*_*EPSP*_(*t*) divided by the amplitude *A* of the parameter in the electrical model, drives the induced slow fluctuation. Consequently, CBF signals simultaneously influence the changes of the cerebral blood volume (CBV) *ν*(*t*) and deoxyhemoglobin content (DOH) *q*(*t*) as described by the following equations:

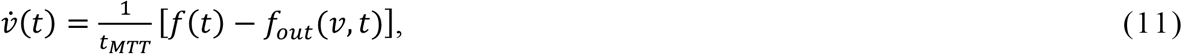

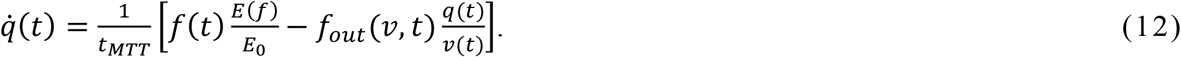

The mean transit time *t*_*MTT*_ scales both differential equations for passing a bolus of the blood through the vein. To estimate CBV changes, Equation 11 models a difference between the blood inflow *f*(*t*) and the blood outflow *f*_*out*_(*ν, t*). Subsequently, we can calculate the changes of DOH using the dynamics of CBF and CBV by regarding oxygen extraction fraction *E*(*f*) in Equation 12. Parameter *E*_0_ is the net oxygen extraction fraction at rest,

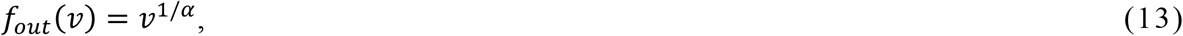

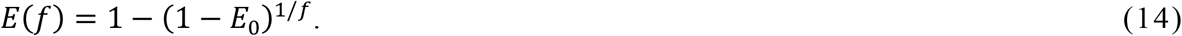

Equation 13 provides the relationship between CBF and CBV, where Grubb *et al*.^2^ empirically found *α* is 0.38. Equation 14 is a non-linear function of CBF, and describes an effect of CBF on the oxygen extraction fraction, see the reference for details.^1^ Using CBV and DOH, we can calculate simulated BOLD signals *y*_*BOLD*_:

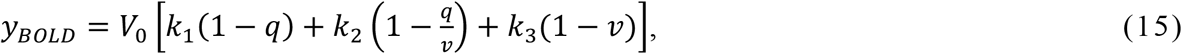

where *V*_0_ is the resting blood volume fraction, and parameters *k*_1_, *k*_2_, and *k*_3_ depend on the magnetic field strength as follows:

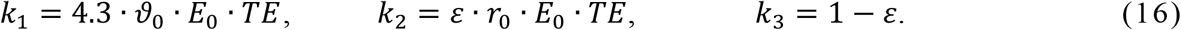

Parameters *ϑ*_0_, *TE, ε*, and *r*_0_ are the frequency offset for 3 T scanner, the echo time, the ratio of intra/extra-vascular signal, and the sensitivity of changes in intra-vascular signal relaxation rate with changes in oxygen saturation, respectively.^1^ The parameter values of the BW model for BOLD signals are given in Table 2.

**S-Figure 1.**
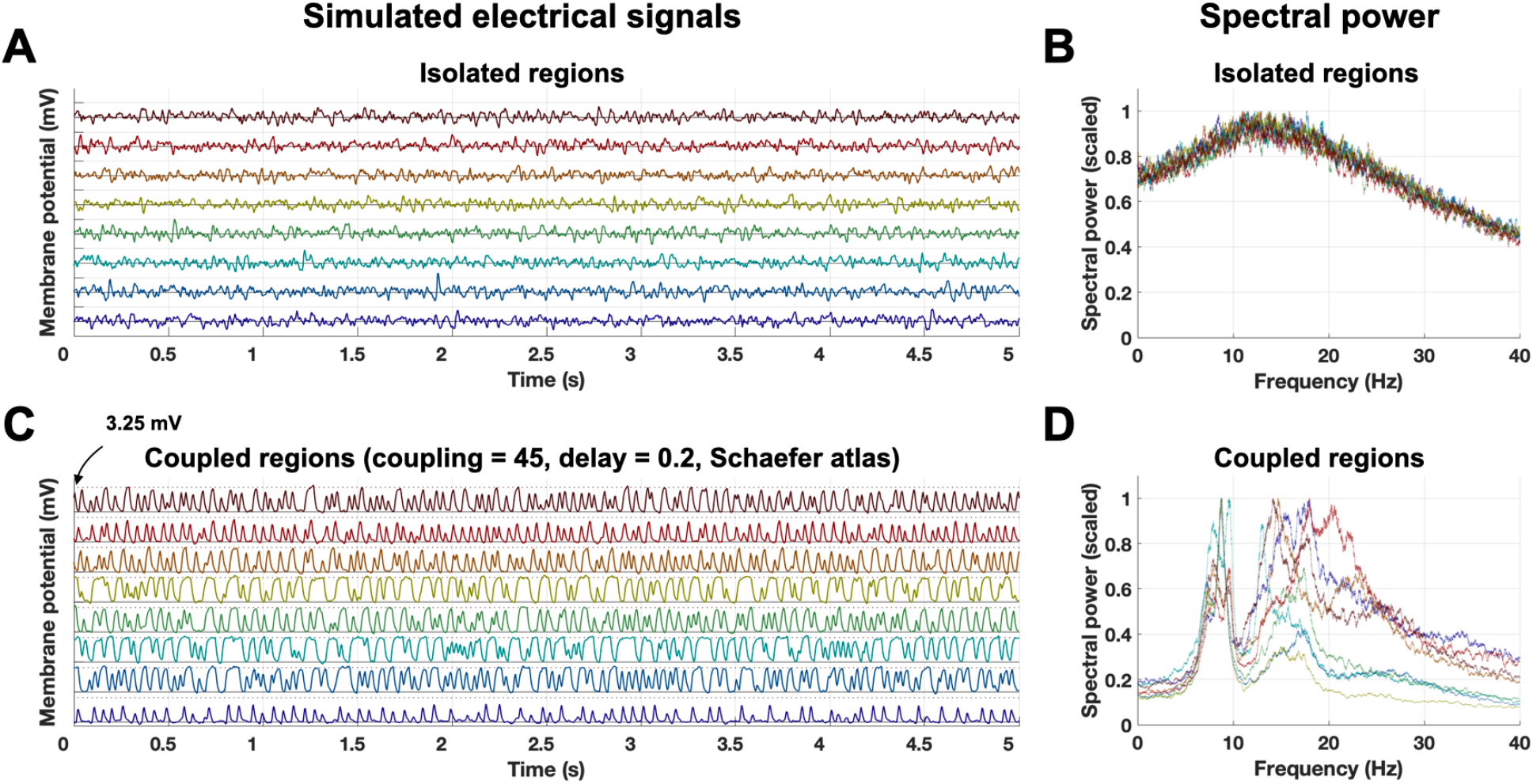
Example of simulated excitatory post-synaptic potentials (EPSP) and their spectral power distributions for isolated (global coupling = 0) and coupled (global coupling = 45 and global delay = 0.2 on the optimal model parameters for the Schaefer atlas) cases. **(A, C)** EPSP signals for **(A)** the isolated regions and for **(C)** the coupled regions. **(B, D)** Spectral power distributions of the EPSP signals in (A, C), respectively. The peaks of the maximal power of the distributions for the isolated regions in (B) are around 13 Hz. The dotted horizontal lines in (C) indicate the maximum EPSP (3.25 mV), which is the specified value as the maximal EPSP kernel in Table 2.

**S-Figure 2.**
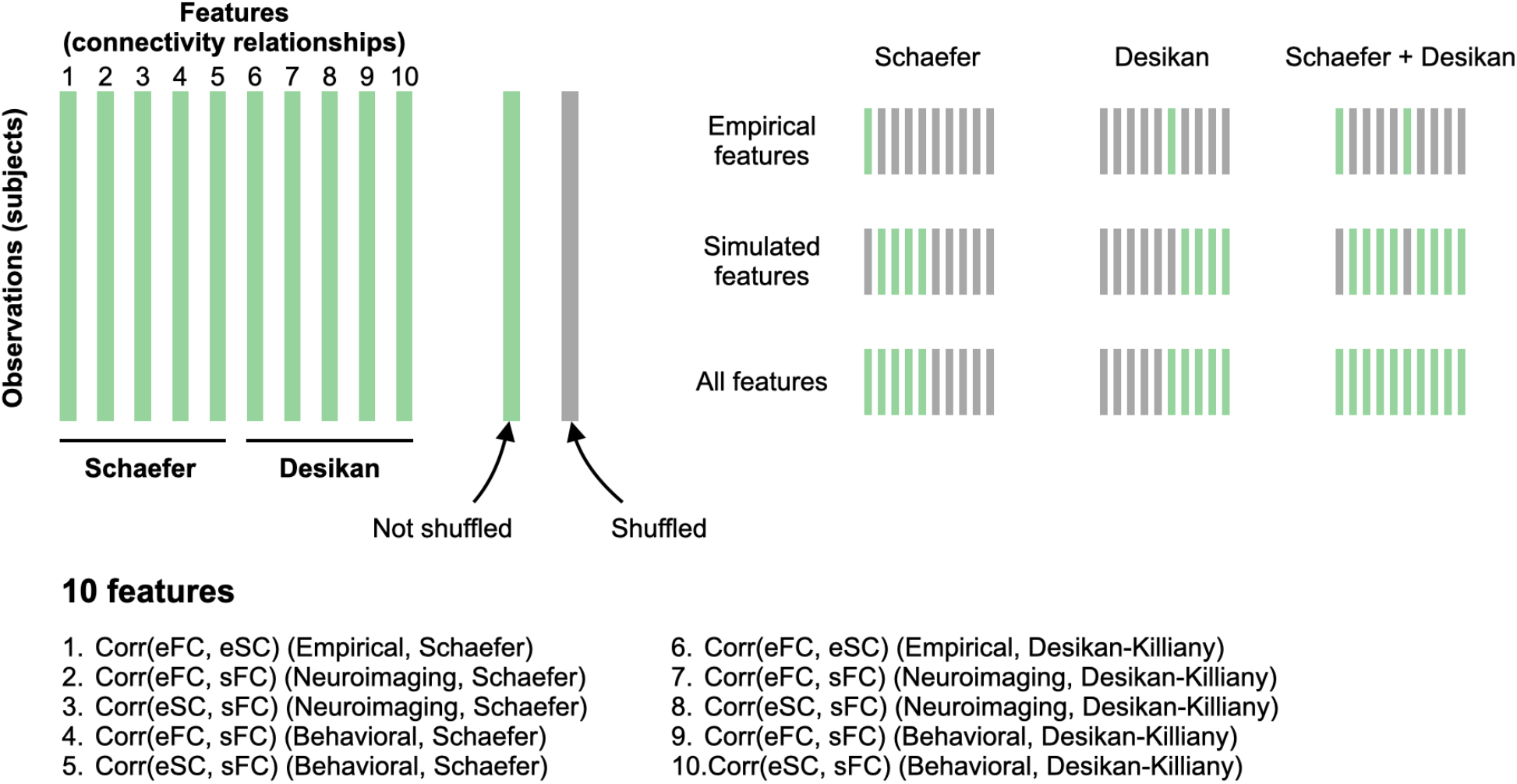
Feature conditions for PD classification. Ten connectivity relationships were used for PD classification as features. To investigate the impact of simulated results on the prediction performance, we selectively shuffled features (gray bars). Shuffling is done in each feature separately, i.e., shuffling within feature which gives the same distribution but randomized feature values across subjects. Abbreviations: FC = functional connectivity; PD = Parkinson’s disease; SC = streamline count.

**S-Figure 3.**
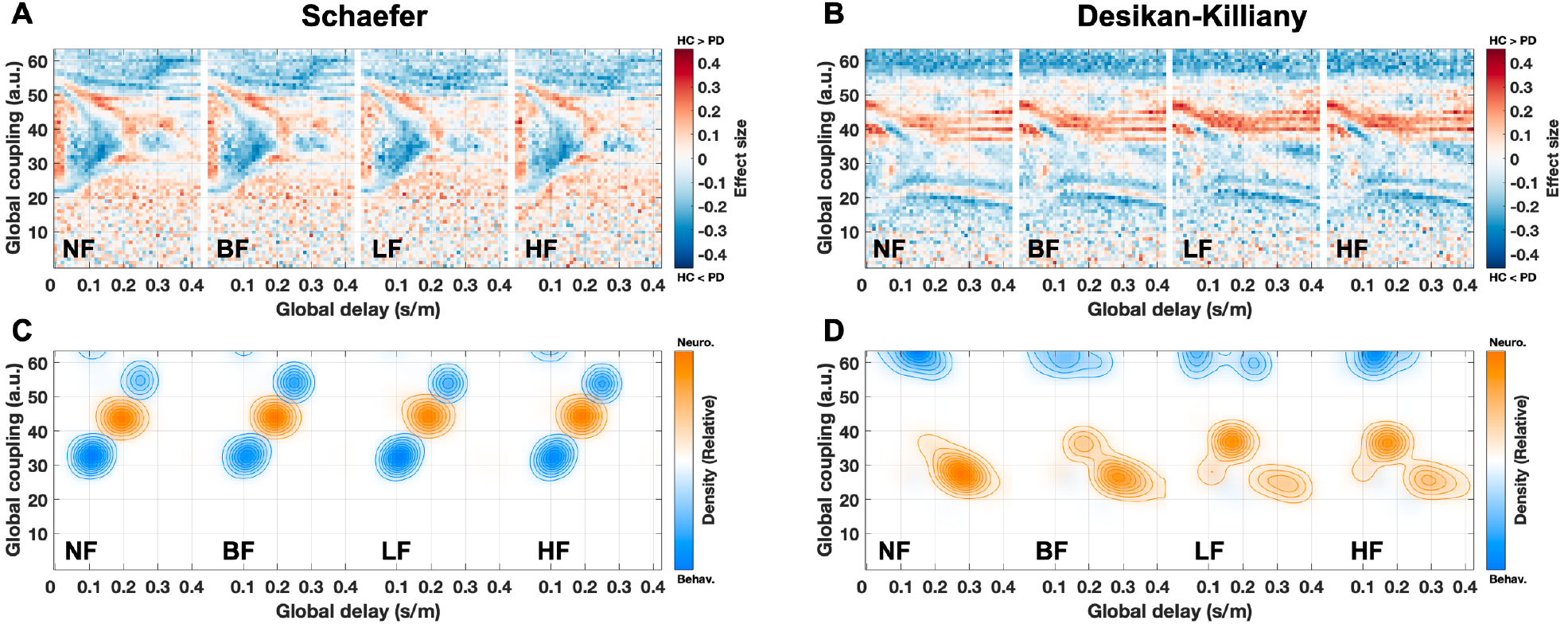
Parameter maps of the effect size of the difference of eSC-sFC correlation values between PD and HC subject groups used for the behavioral model fitting. The filtering conditions are indicated in the plots for **(A)** the Schaefer atlas and **(B)** the Desikan-Killiany atlas. Effect sizes in the (*τ*_*global*_, *C*)-parameter plane were calculated by a non-parametric Wilcoxon rank-sum two-tailed test between PD and HC in the eSC-sFC correlation values for each parameter point. **(C, D)** Distributions of optimal parameters derived from the neuroimaging model fitting (orange, all subjects, n=116) and the behavioral model fitting (blue, repeated sub-sampling, n=1000) for **(C)** the Schaefer atlas and **(D)** the Desikan-Killiany atlas. Abbreviations: PD = Parkinson’s disease; HC = healthy controls; NF = no filtering; BF = broad band ([0.01,0.1] Hz); LF = low-frequency band ([0.01,0.05] Hz; HF = high-frequency band ([0.05,0.1] Hz).

**S-Figure 4.**
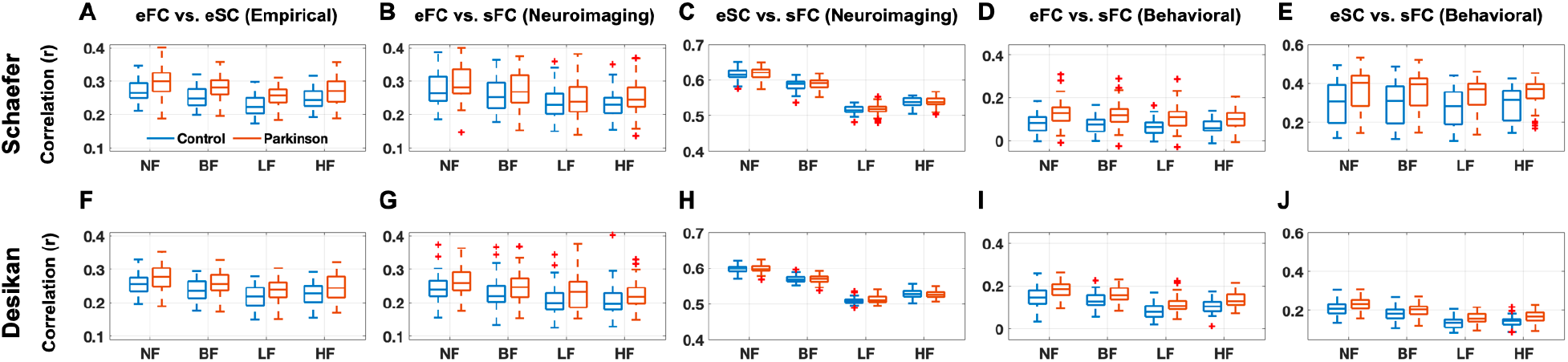
Comparison of connectivity correspondences between PD and HC subjects as reflected by the connectivity relationships of empirical and simulated results for **(A-E)** the Schaefer atlas and **(F-J)** the Desikan-Killiany atlas for **(A, F)** the empirical structure-function relationship (eFC vs. eSC), **(B, C, G, H)** functional (eFC vs. sFC) and structure-function (eSC vs. sFC) relationships for the neuroimaging model fitting, and **(D, E, I, J)** connectome relationships (eFC vs. sFC and eSC vs. sFC) for the behavioral model fitting. Abbreviations: NF = no filtering; BF = broad band ([0.01,0.1] Hz); LF = low-frequency band ([0.01,0.05] Hz; HF = high-frequency band ([0.05,0.1] Hz); FC = functional connectivity; SC = streamline count.

**S-Figure 5.**
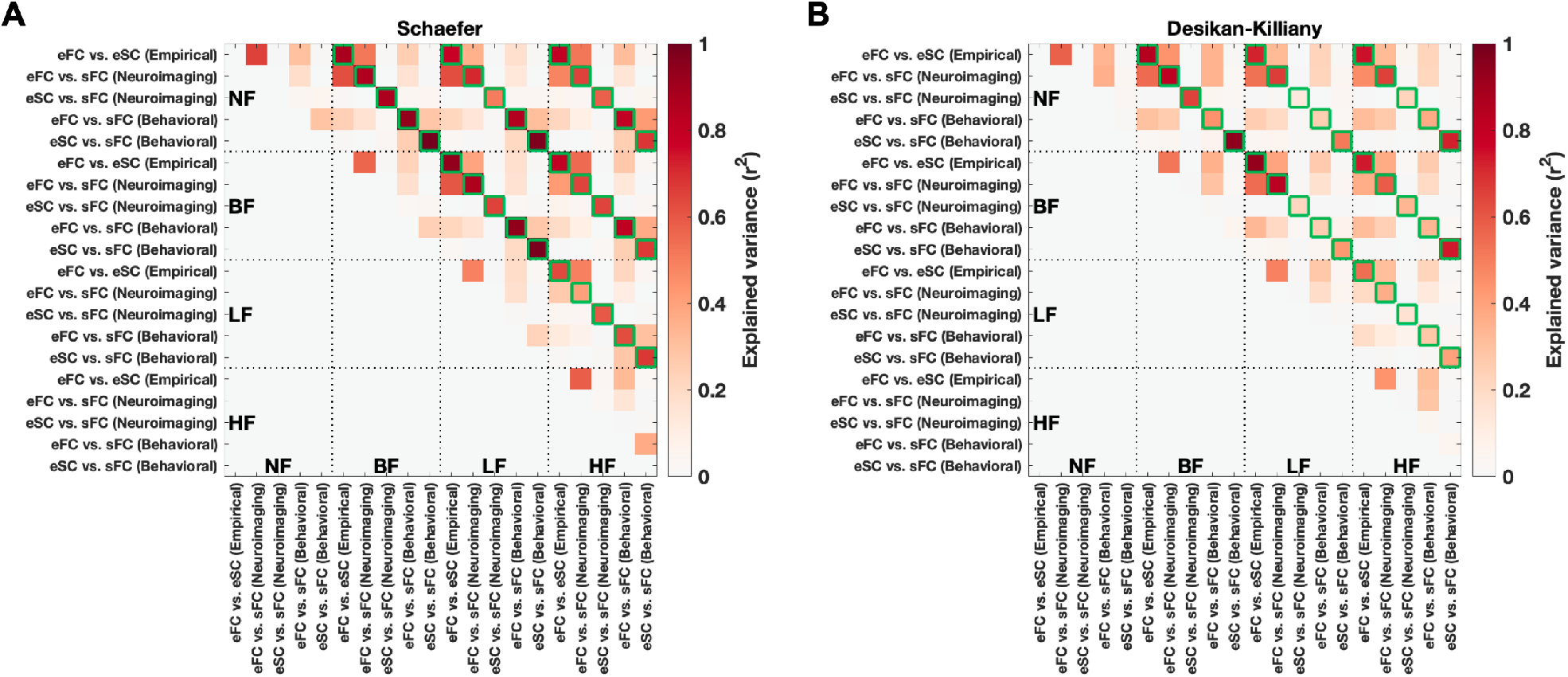
Explained variances (EV) between five connectivity relationships in intra-/inter-conditions for **(A)** the Schaefer atlas and **(B)** the Desikan-Killiany atlas. The five connectivity relationships are corr(eSC, eFC) (empirical), corr(eFC, sFC) (neuroimaging), corr(eSC, sFC) (neuroimaging), corr(eFC, sFC) (behavioral), and corr(eSC, sFC) (behavioral). Due to the four considered temporal filtering conditions, the intra-/inter-EVs were obtained using 20 connectivity relationships (see the axes). The green boxes are for the same types of connectivity relationships under different filtering conditions. Abbreviations: NF = no filtering; BF = broad band ([0.01,0.1] Hz); LF = low-frequency band ([0.01,0.05] Hz; HF = high-frequency band ([0.05,0.1] Hz).

**S-Figure 6.**
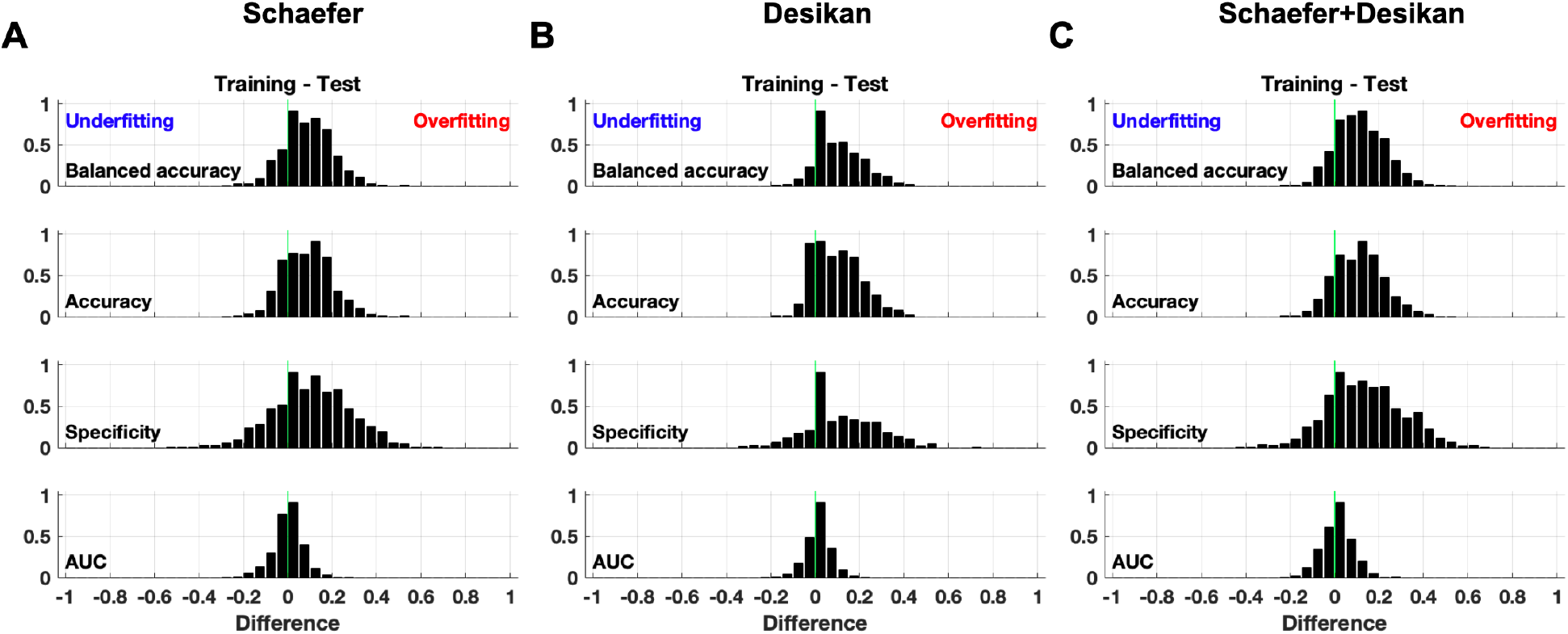
Differences of model performance between training and test sets (Training – Test) for PD prediction including all filtering conditions and all features for the **(A)** Schaefer atlas, **(B)** Desikan-Killiany atlas, and **(C)** multiple atlases, i.e., the Schaefer and Desikan-Killiany atlases. The green vertical lines indicate zero differences. The positive differences are overfitting cases, and the negative ones are underfitting. Abbreviation: AUC = area-under-curve.

**S-Figure 7.**
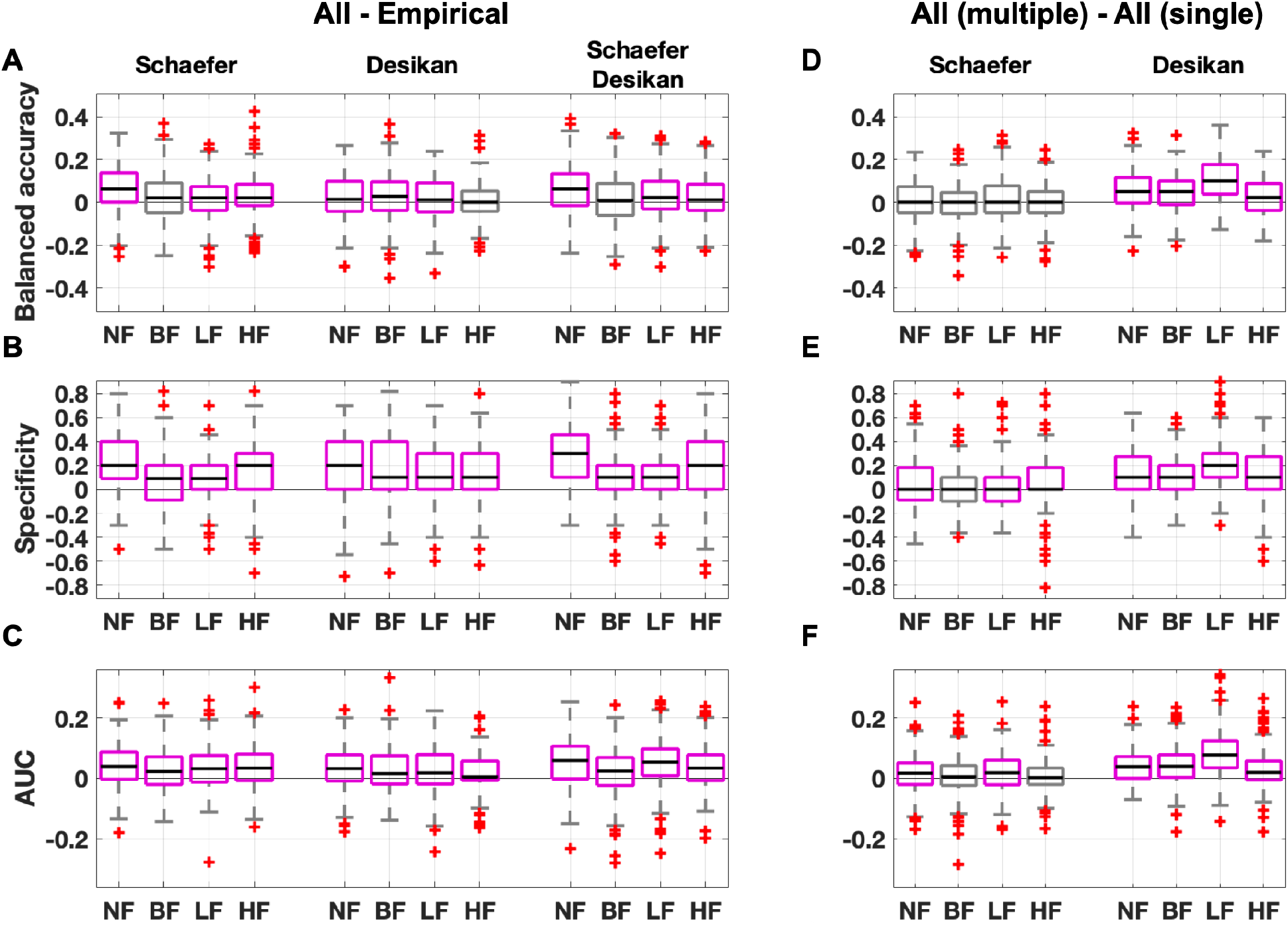
Comparisons of prediction performance by **(A, B, C)** differences (All - Empirical) for the Schaefer, Desikan-Killiany, and multiple (Schaefer and Desikan-Killiany) atlases for **(A)** balanced accuracy, **(B)** specificity, and **(C)** AUC of ROC curves from different feature conditions for each filtering condition. **(D, E, F)** Differences by ‘All features’ (multiple atlases - a single atlas) for **(D)** balanced accuracy, **(E)** specificity, and **(E)** AUC of ROC curves. The purple boxes indicate significantly different conditions (Wilcoxon signed-rank two-tail test and Bonferroni corrected *p* < .05). Abbreviations: NF = no filtering; BF = broad band ([0.01,0.1] Hz); LF = low-frequency band ([0.01,0.05] Hz; HF = high-frequency band ([0.05,0.1] Hz).

**S-Figure 8.**
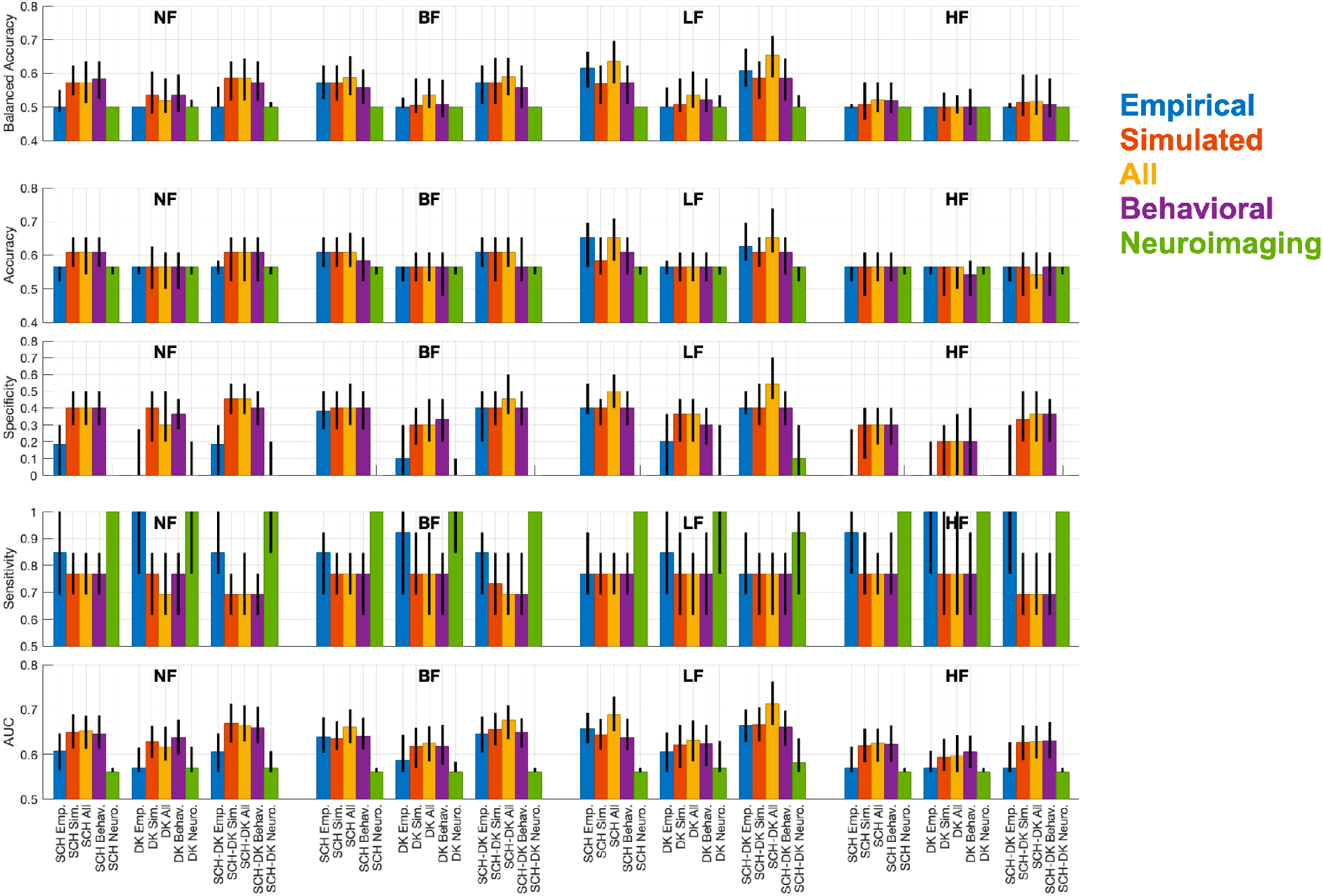
Prediction performances as given by the balanced accuracy, accuracy, specificity, sensitivity, and AUC of ROC curves using optimal simulated connectomes from the behavioral fitting only (purple) and from the neuroimaging fitting only (green) as additional feature cases to those presented in Fig. 6 in the main text (also the same here for comparison). The error bars indicate interquartile ranges, and the heights of bars are medians. Abbreviations: NF = no filtering; BF = broad band ([0.01,0.1] Hz); LF = low-frequency band ([0.01,0.05] Hz; HF = high-frequency band ([0.05,0.1] Hz); SCH = Schaefer; DK = Desikan-Killiany.

**S-Figure 9.**
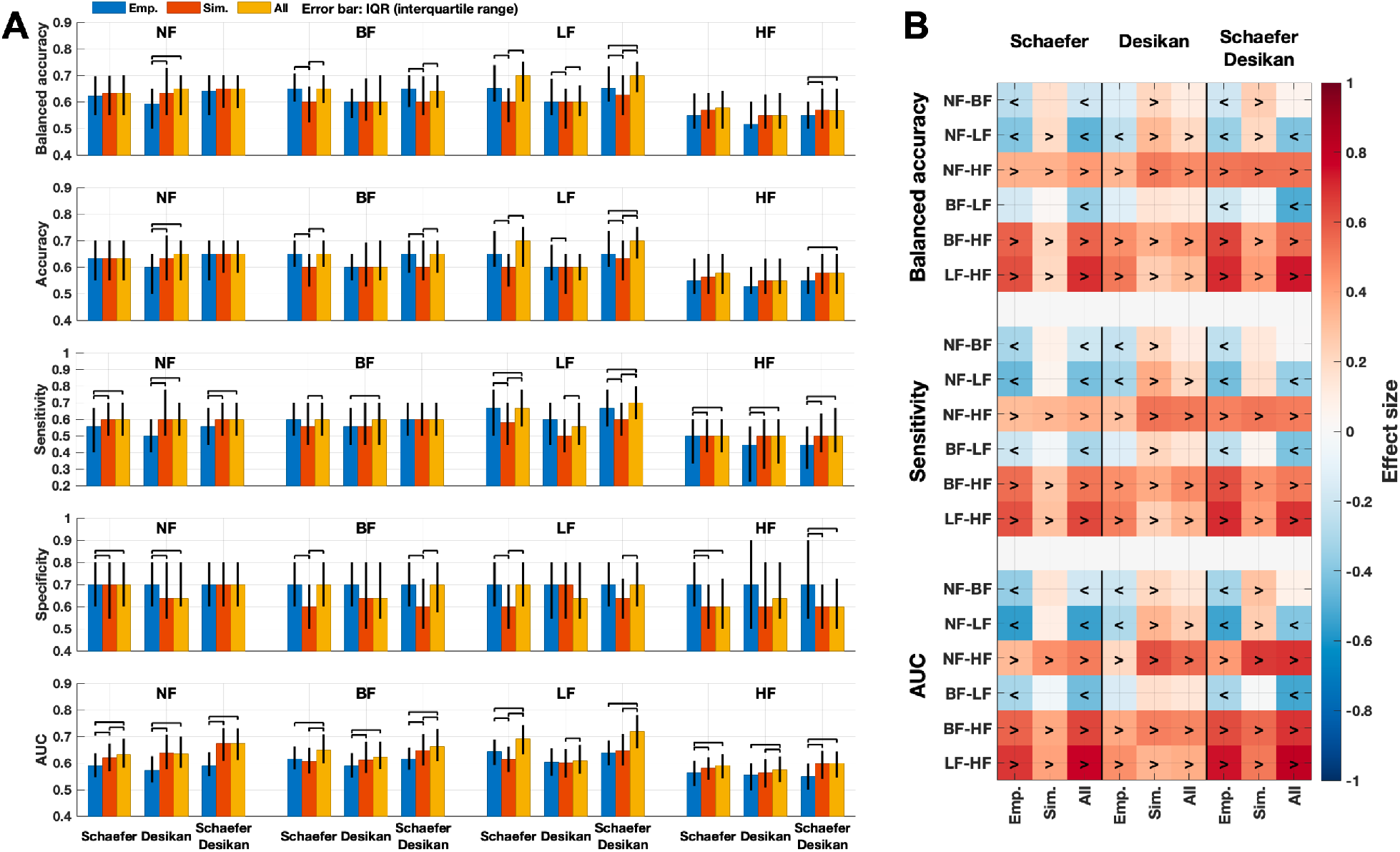
Summary of the performance of PD classification using the three different feature conditions: empirical features (blue bars), simulated features (red bars), and all features (yellow bars) after incorporating the age controlling and the behavioral model fitting during the nested cross-validation (Fig. 2 in Methods) for the balanced subject configuration (n=99, S-Table 1). **(A)** Median values of the balanced accuracy, accuracy, sensitivity, specificity and area-under-curve (AUC) of the receiver operating characteristics (ROC) curves for all considered parcellations and filtering conditions are shown in each panel. The error bars indicate the interquartile range across iterations of the outer loop of the nested cross-validation procedure (Fig. 2). The black lines connecting two conditions indicate significantly different performance between feature conditions. **(B)** Effect sizes between filtering conditions for each feature condition. The signs ‘<’ and ‘>’ indicate which condition is significantly larger than the other. For example, ‘<’ sign for ‘NF-LF’ indicated on the vertical axes means NF < LF for a given performance indicated on the horizontal axes. The Wilcoxon signed-rank two-tail test was used for comparisons across feature and filtering conditions (Bonferroni corrected statistics). Abbreviations: PD = Parkinson’s disease; NF = no filtering; BF = broad band ([0.01,0.1] Hz); LF = low-frequency band ([0.01,0.05] Hz; HF = high-frequency band ([0.05,0.1] Hz).

**S-Table 1.** Demography of a balanced subject configuration (excluding 17 oldest patients from 116 subjects).

| Groups | Mean (standard deviation) years |  | Statistical tests | p-values |
| --- | --- | --- | --- | --- |
|  | **All subjects** |  | **Chi-square goodness-of-fit test** |  |
| All | 56.62 (9.24) |  | 99 subjects | 0.235 |
|  | **Healthy controls** | **Patients** | **Wilcoxon rank-sum two-tail test** |  |
| All | 55.02 (9.69) | 58.31 (8.42) | 51 healthy controls versus 48 patients | 0.062 |
| Female | 56.52 (9.40) | 60.80 (8.96) | 21 healthy controls versus 20 patients | 0.201 |
| Male | 53.97 (9.74) | 56.54 (7.53) | 30 healthy controls versus 28 patients | 0.156 |
|  | **Females** | **Males** | **Wilcoxon rank-sum two-tail test** |  |
| All | 58.61 (9.43) | 55.21 (8.84) | 41 females versus 58 males | 0.095 |

